# Divergent patterns of richness and density in the global soil seed bank

**DOI:** 10.1101/2023.11.08.566230

**Authors:** Alistair G. Auffret, Emma Ladouceur, Natalie S. Haussmann, Petr Keil, Eirini Daouti, Tatiana G. Elumeeva, Ineta Kačergytė, Jonas Knape, Dorota Kotowska, Matthew Low, Vladimir G. Onipchenko, Matthieu Paquet, Diana Rubene, Jan Plue

**Affiliations:** Department of Ecology, Swedish University of Agricultural Sciences, 75 007 Uppsala, Sweden; Department of Biology, University of Prince Edward Island, PE, Canada; Canadian Centre for Climate Change and Adaptation, University of Prince Edward Island, St. Peter’s Bay, PE, C0A 2A0, Canada; School of Climate Change and Adaptation, University of Prince Edward Island, Charlottetown, PE, C1A 4P3, Canada; German Centre for Integrative Biodiversity Research (iDiv) Leipzig-Halle-Jena, Puschstraße 4, 04103 Leipzig, Germany; Department of Geography, Geoinformatics and Meteorology, University of Pretoria, Lynnwood Rd, Hatfield 0002, South Africa; Faculty of Environmental Sciences, Czech University of Life Sciences Prague, Kamýcká 129, 165 00 Praha-Suchdol, Czech Republic; Department of Crop Production Ecology, Swedish University of Agricultural Sciences, 75 007 Uppsala, Sweden; Moscow Lomonosov State University, Moscow, 119234, Russia; Institute of Nature Conservation, Polish Academy of Sciences, Kraków, Poland; ‘Lendület’ Landscape and Conservation Ecology Group, Institute of Ecology and Botany, MTA–HUN-REN Centre for Ecological Research, Vácrátót, Hungary; Theoretical and Experimental Ecology Station (SETE), National Center for Scientific Research (CNRS), Moulis, France; Greensway AB, Gerda Nilssons väg 2, 756 51 Uppsala, Sweden; Swedish Biodiversity Centre, Swedish University of Agricultural Sciences, 75 007 Uppsala, Sweden

## Abstract

Soil seed banks are an important component of plant population and community dynamics, buffering temporal heterogeneity and allowing populations to recover following disturbance. At the same time, patterns in soil seed banks are likely to reflect scale-dependent patterns in above-ground vegetation, patterns that differ strongly across world regions. Here, we investigate components of diversity in the soil seed bank across global biomes and ecosystems. We found that patterns of species richness in the soil seed bank diverged from patterns of seed density, although there were high levels of uncertainty, especially at smaller spatial scales. Importantly, habitat degradation consistently led to lower richness, potentially limiting future ecosystem recovery. Among ecosystems, mediterranean and tropical regions had high species richness, but seed density m^−2^ in the soil was highest in wetlands and arable systems. Lower seed densities were found in both high-diversity tropical biomes that are characterised by short-lived seeds, and low-diversity boreal and tundra biomes with more stable established vegetation. Our findings of divergences between species richness and seed density across global biomes, and how they are impacted by habitat degradation, give valuable insights to the understanding of plant biodiversity worldwide.

## Introduction

In an era of global environmental change, the persistence of populations in newly-unsuitable conditions and the establishment of new populations following disturbance are key in determining changes in biodiversity over time and in space. For vascular plants, the soil seed bank can play a vital role in both processes. Soil seed banks are the communities of persistent seeds being naturally stored in the soil that contribute to vegetation regeneration after disturbance^1,2^. Plant species that form seed banks do so as part of a life-history strategy that allows for the buffering of temporal environmental heterogeneity^3–5^, and as such, seed banks are an important component for understanding plant (meta)population and community dynamics^6,7^.

Despite the potential advantage of producing seeds that persist in the soil, not all species contribute to soil seed banks. Instead, the ability to form seed banks is a functional strategy for plants to (re)establish and maintain populations, which has evolved in response to both predictable and unpredictable disturbance regimes^8^. The species that do form seed banks also vary in terms of the number of seeds produced, and the longevity of seeds in the soil^9^. The disturbances that have contributed to the development of seed banking are not randomly distributed in space, and as such plant communities in different bioclimatic environments should rely on seed banking to different extents, with implications for local seed bank size and composition. For example, in arid systems dominated by drought, it is advantageous to have an annual life history and form persistent seed banks to buffer annual changes in water availability^10,11^. A similar pattern exists in temperate Europe, where species with warmer distributions have been found to have a higher seed bank persistence than more northerly species^12^. In tropical systems, where climatic conditions are broadly stable and exposure to freezing temperatures and drought conditions is rare, relatively few species produce seeds that are able to survive desiccation^13,14^ – a trait associated with long-term persistence – which may result in relatively small and species-poor soil seed banks.

While it can be interesting to broadly compare plant communities from different world regions, ecological assemblages can differ considerably among habitats within the same region. The frequency of disturbance within a habitat is generally mirrored by the strategies of the resulting plant communities^15^. Therefore, different habitats could be expected to support different soil seed banks, in terms of the number of species and the density of seeds present. For example, in temperate and boreal regions, forests are dominated by long-lived trees, with the understorey often covered by species that largely rely on vegetative reproduction rather than seed banking^16,17^. Species-rich grazed grasslands in the same regions are characterised by regular, low-intensity and spatially-heterogeneous disturbances, and as such harbour plants with a wide range of regeneration strategies^18,19^. Intensively-used arable lands on the other hand are well-known to support large communities of ruderal species with annual life-histories and high seed production^20,21^. As such, habitat is an important component in understanding how different functional traits are represented in different regions, while differences between broad habitat types can differ in different world regions^22^. Moreover, communities in the same broad habitat categories can differ in their responses to environmental degradation in different regions^23^. Therefore, when studying plant communities at global scales, it is important to consider the potential roles of different habitats and environmental degradation driving any variation within or between regions. This could be especially relevant in the case of soil seed banks, which exist partially as a strategy to respond to disturbance.

Studying patterns of soil seed bank diversity across large spatial scales and geographic gradients can be useful for understanding the biogeography and macroecology of their role in biodiversity maintenance and recovery^24–26^. However, biodiversity is known to be scale-dependent, and comparisons of soil seed banks across ecoregions and habitats are therefore likely to be influenced by the scale at which diversity is estimated. It has been established that the number of species in soil seed banks increases with area^24,25^, but while it is well-known that this accumulation of species varies according to habitat and region in the established vegetation as well as in other taxa^27–30^, such patterns are still largely unknown in the soil seed bank, at least not at spatial scales that span multiple biomes.

Integrating a scale-explicit approach to understanding diversity in the soil seed bank across habitats, within and between ecoregions and in response to habitat degradation is an important step for plant community and macroecology, allowing critical insights for both broad scale patterns of biodiversity and the context-dependent interpretations of local dynamics.

While individual seed bank studies are usually (through necessity) small-scale comparisons across limited environmental gradients within a specific region, global analysis allows the large-scale comparison of biodiversity of the seed bank. Here, we use a global database of soil seed banks, harnessing 1442 studies from all seven of the world’s continents (Fig. 1), to reveal how broad differences in climatic conditions across world regions, and differences in disturbance regimes across habitat types, combine to determine patterns in both the richness and density of soil seed bank communities at different spatial scales. The global seed bank database^31^ is the result of a comprehensive search of the seed bank literature, providing 3096 records containing information regarding the richness and/or density of the soil seed bank at a given location (Methods). To investigate patterns of seed bank diversity among ecosystems and global biomes, we used both the WWF map of global ecoregions^32^ and the habitat descriptions used by the authors of the component studies to separate records into 15 ecosystems. These ecosystems are defined as the combination of biome (e.g. forest, grassland) and ecoregion (e.g. temperate, tropical). Each ecosystem was assigned to either the terrestrial, aquatic or transitional realm^33^; see Fig. 1 and Table 1.

**Figure 1.**
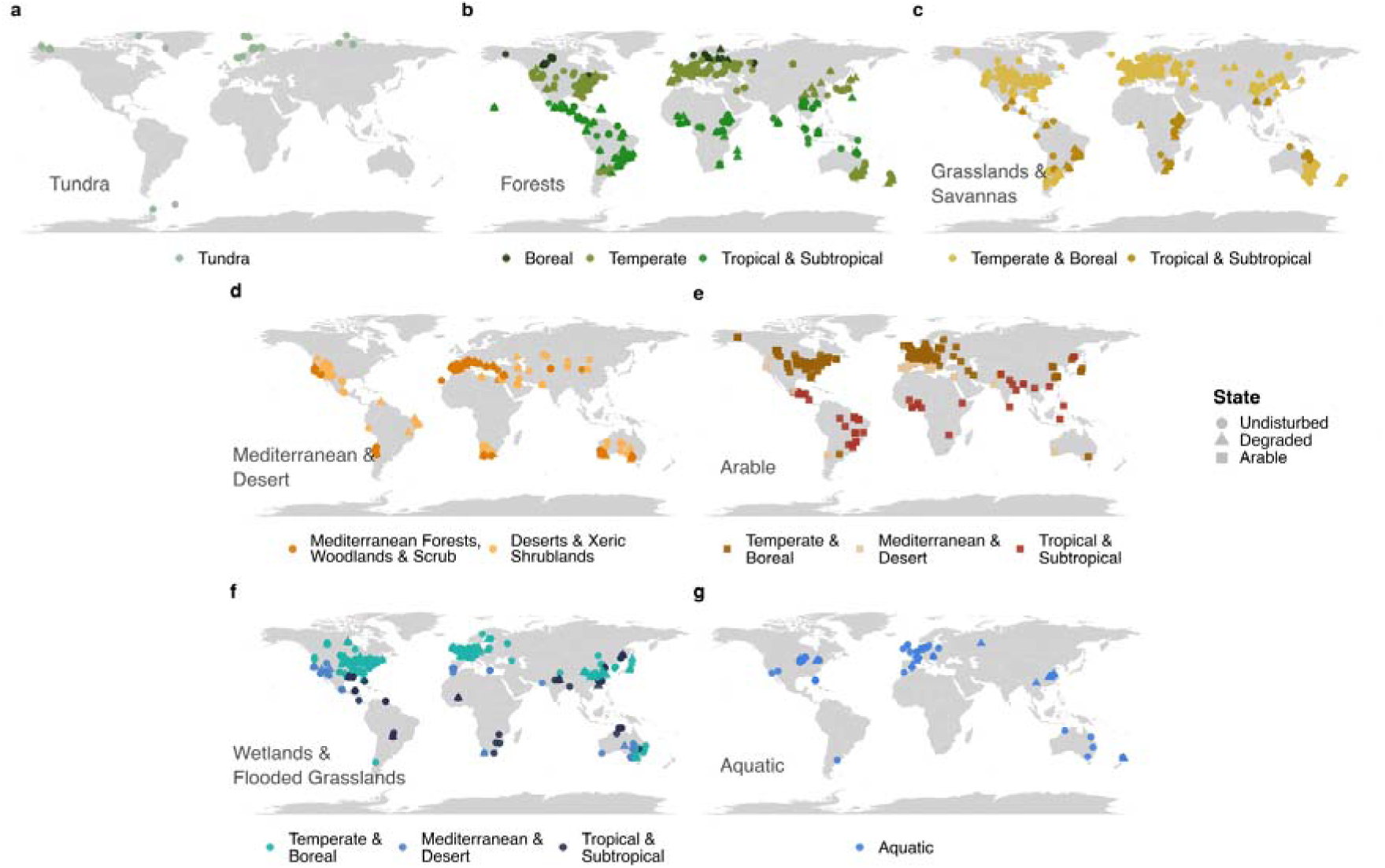
The global record of the soil seed bank. Points indicate the location of each record collected from the literature (n=3096; some records have the same location). Records are first split according to biome (e.g. tundra, forests; panels a-g), with each biome split according to one or more ecoregions (e.g. boreal, tropical; colours of points). The combination of biome and ecoregion results in 15 unique ecosystems. Each record is also designated either as occurring in undisturbed or degraded ecosystems, apart from arable records which have their own category (shapes of points). We also categorise each biome as belonging to the **a-e** Terrestrial, **f** Transitional, or **g** Aquatic realm. See Methods for a more detailed description of all categories.

**Table 1.**
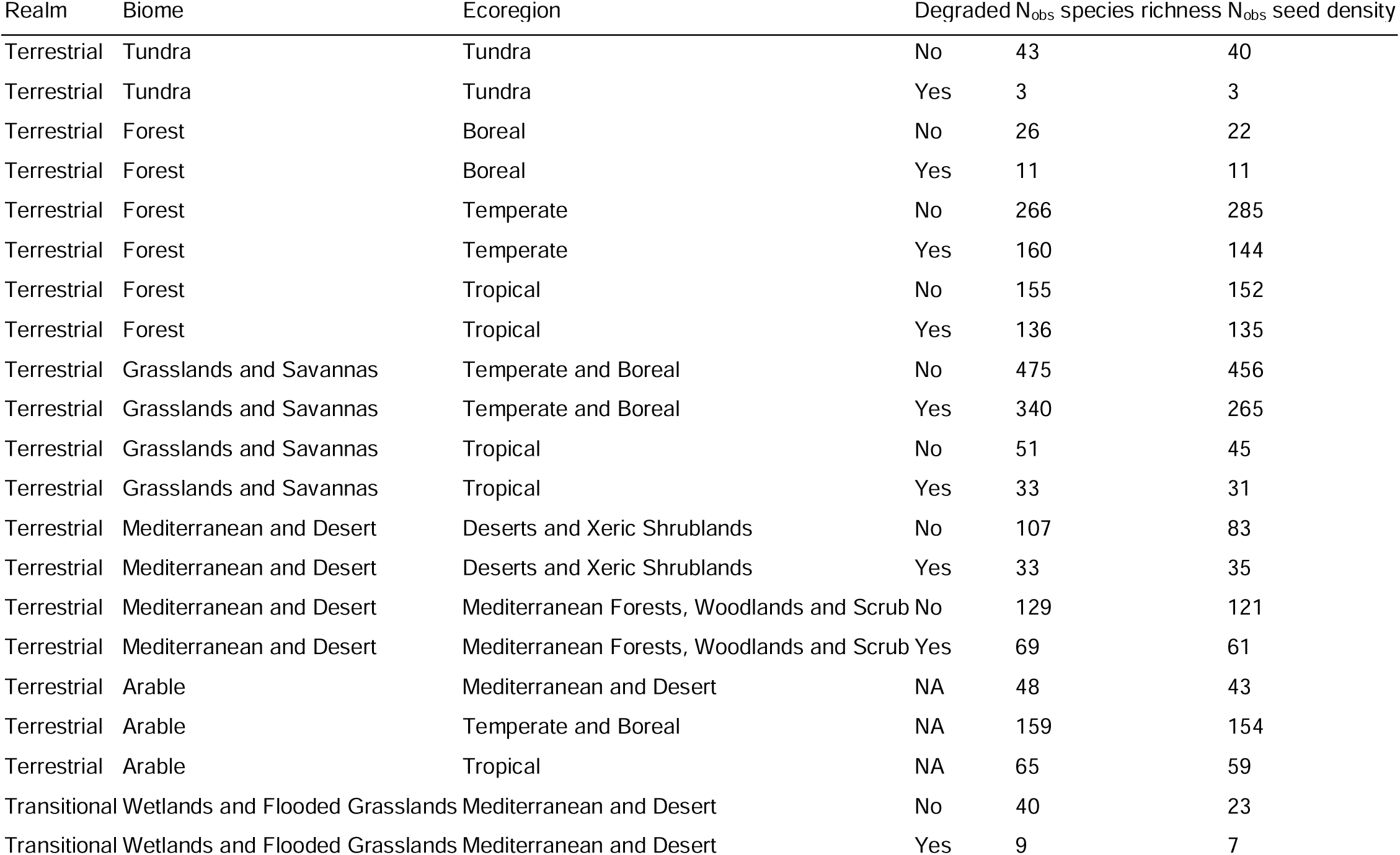
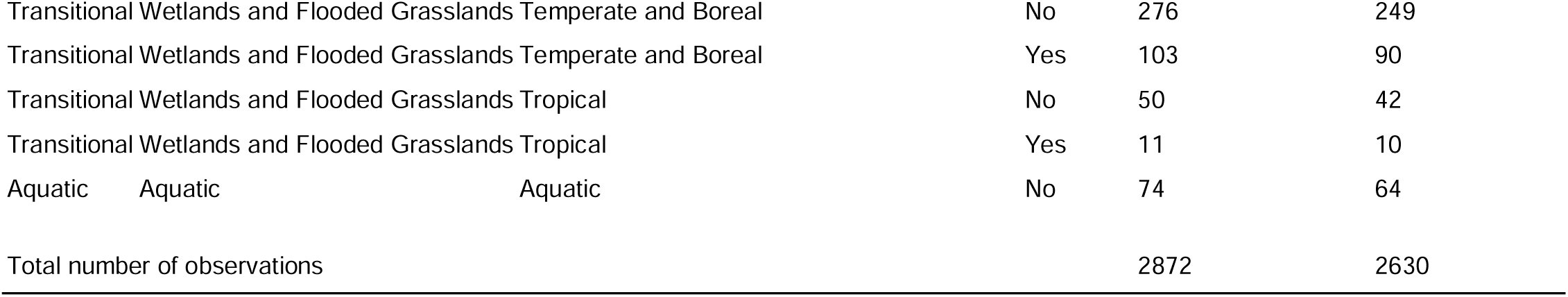
Division of soil seed bank observation into ecosystems. Biome and ecoregion are defined according to author-reported habitat types and geographical association to global ecoregions (Methods). The combination of biome and ecoregion is what we refer to as an ecosystem. ‘Degraded’ indicates whether the area samples were taken from was considered to be degraded by authors of the original study. The number of observations (N_obs_) collected for each ecoregion, and degraded status listed for species richness, and seed density m^−2^.

We used a hierarchical Bayesian modelling approach to investigate different components of the soil seed bank separately per biome, with ecoregion, habitat degradation (binary: undisturbed or degraded) and sampling effort (area sampled and number of sites) included as predictor variables. Specifically, we investigated [1] species richness, finding highest richness in tropical forests and mediterranean ecosystems, but differences among ecosystems and biomes, reflecting our broad knowledge of global patterns in the aboveground vegetation; [2] seed density (m^−2^), finding highest values in wetlands and arable ecosystems; and [3] the impact of habitat degradation, finding consistently lower richness in degraded habitats, with patterns of density. Importantly, we found that patterns of species richness in the soil seed bank diverged from patterns of density. That is, a high density of seeds in the soil seed bank does not necessarily translate to a large number of species, and that this varies among ecosystems, biomes and in response to degradation.

## Results

The global seed bank database^31^ contains the results of studies carried out in 94 countries across the world, including observations of more than 35 million seeds using a large range in sampling effort and extent (for a more in-depth summary of the data, see Methods). In such a large and heterogeneous dataset, there was generally a large variation in seed bank richness and density within and between ecosystems. Here, we describe noteworthy results in terms of overlap in 50% and 90% credible intervals (hereafter CIs) and consistency in the relative values of model estimates that are rounded to whole numbers (i.e. whole species or whole seeds m^−2^). All model results and further supplementary information can be found in Tables S1-S4, and Figures S2a-g, and full details of statistical models, including the definitions, interactions, inclusion and transformation of and between predictor variables are described in the Methods.

### Seed bank richness and density across ecosystems and biomes

Using models analysing species richness to predict average soil seed bank diversity for each ecosystem and at multiple spatial scales, we found that predicted species richness generally increased from the 0.01m^2^ alpha scale to the 15 m^2^ gamma scale. Estimated richness was always higher at the gamma scale, with 50% credible intervals not overlapping in all undisturbed and 80% of degraded (including arable) systems. Relative values of predicted richness varied according to scale, with one example being that boreal and temperate grasslands had a higher predicted richness than tropical forests at the alpha scale (but with overlapping CIs), while tropical forests had higher richness at the gamma scale (no overlap of 50% CIs). Generally, differences in estimated richness across ecosystems were more clear at the gamma scale, so we place most focus there. In undisturbed systems, mediterranean forests and shrublands were predicted to have the highest richness compared to all eleven other undisturbed ecosystems (no overlap of 50% CIs, overlap with six undisturbed ecosystems at 90% CIs; Figure 2, Table S4). Tundra and aquatic systems generally had the lowest species richness, each having a lower estimate than the other ten undisturbed ecosystems, and lower than eight of the other ecosystems (50% CIs).

**Figure 2:**
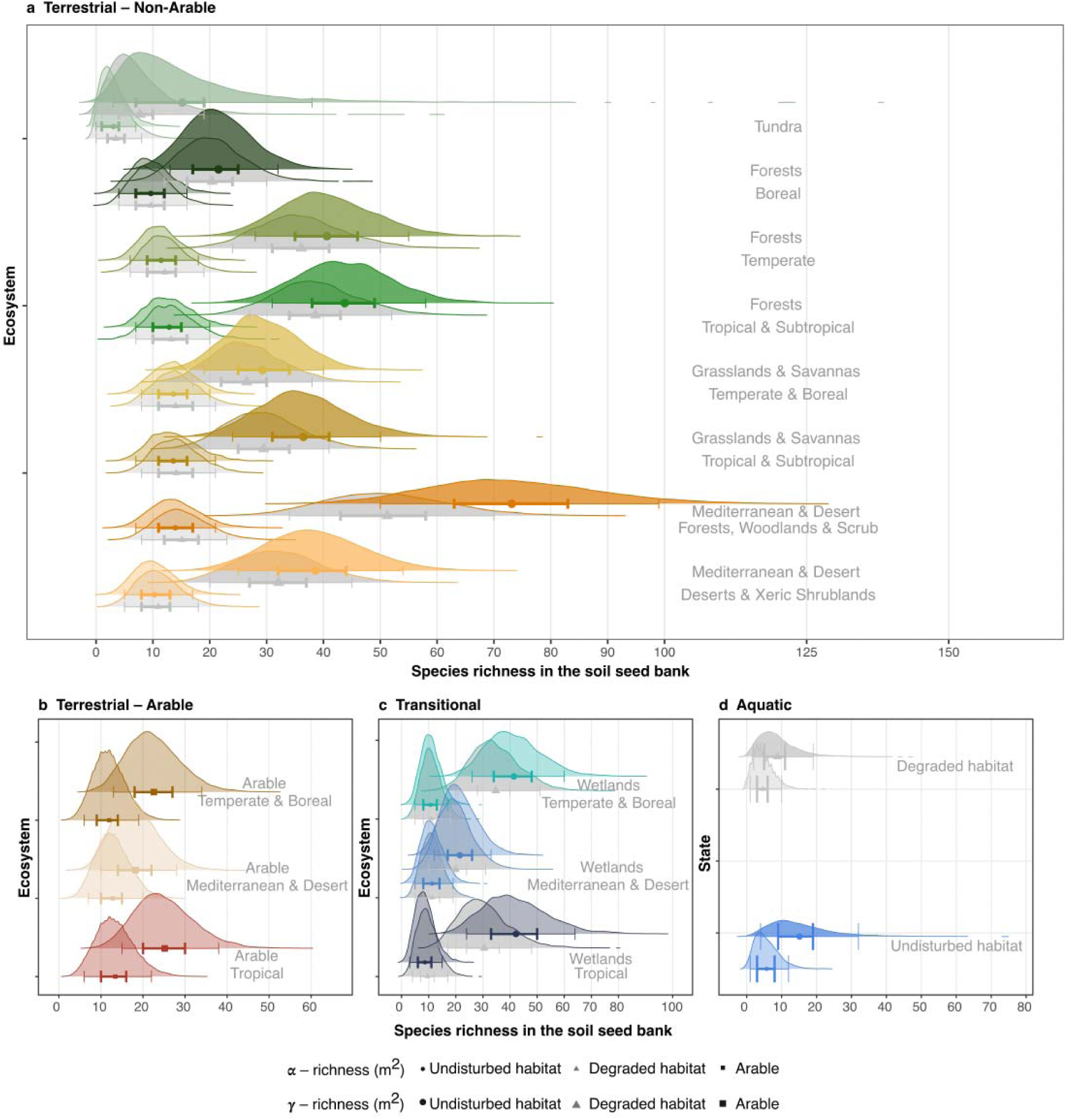
Scale-dependent richness in soil seed banks across ecosystems. Panels **a-d** represent realms, with the terrestrial realm separated into arable and non-arable systems. Each ecosystem is represented by four density curves and point-and-whiskers, showing species richness at the alpha ( ; 0.01 m^2^; lower curves, smaller points) and gamma ( ; 15 m^2^; upper curves, larger points) scales, and in undisturbed (coloured curves and circular points) and degraded (grey curves and triangular points) ecosystems. Note that arable systems are not separated into disturbed and undisturbed and have only two curves per ecosystem and a square point. Points represent average predicted species, surrounded by 50% (thick) and 90% (thin) credible intervals. Density curves represent 24 000 predictive draws of the posterior distributions of overall richness at each scale, within each ecosystem and degradation group.

Undisturbed tropical ecosystems were estimated to be broadly more species-rich than their equivalent temperate and boreal ecosystems. Estimated richness was higher for tropical forests than both boreal and temperate forests, for tropical grasslands compared to temperate and boreal grasslands and tropical arable compared to temperate and boreal arable, while tropical wetlands had similar estimated richness as temperate and boreal wetlands. Despite this consistency, estimates were only clearly different in forest ecosystems (50% CIs), although tropical wetlands were more species-rich than mediterranean and desert wetlands.

The average estimated number of seeds m^−2^ in the soil seed bank was 4 214 m^−2^ (median 3 590) in the (non-arable) terrestrial realm, 9 434 m^−2^ (median 11 001) for arable systems, 10 318 m^−2^ (median 10 802 m^−2^) in the transitional realm, and 8 124 m^−2^ (median 9 562) in the aquatic realm (Figure 3, Table S5). Wetland ecosystems exhibited high seed densities, often higher than other ecosystems within their equivalent ecoregion (90% CIs). For example, tropical wetlands had higher seed densities than tropical forests, and temperate and boreal wetlands had higher seed densities than forests and grasslands in the same regions. Within the terrestrial realm, undisturbed temperate and boreal grasslands had the highest predicted seed density, higher than boreal forests and temperate forests (90% CIs).

**Figure 3:**
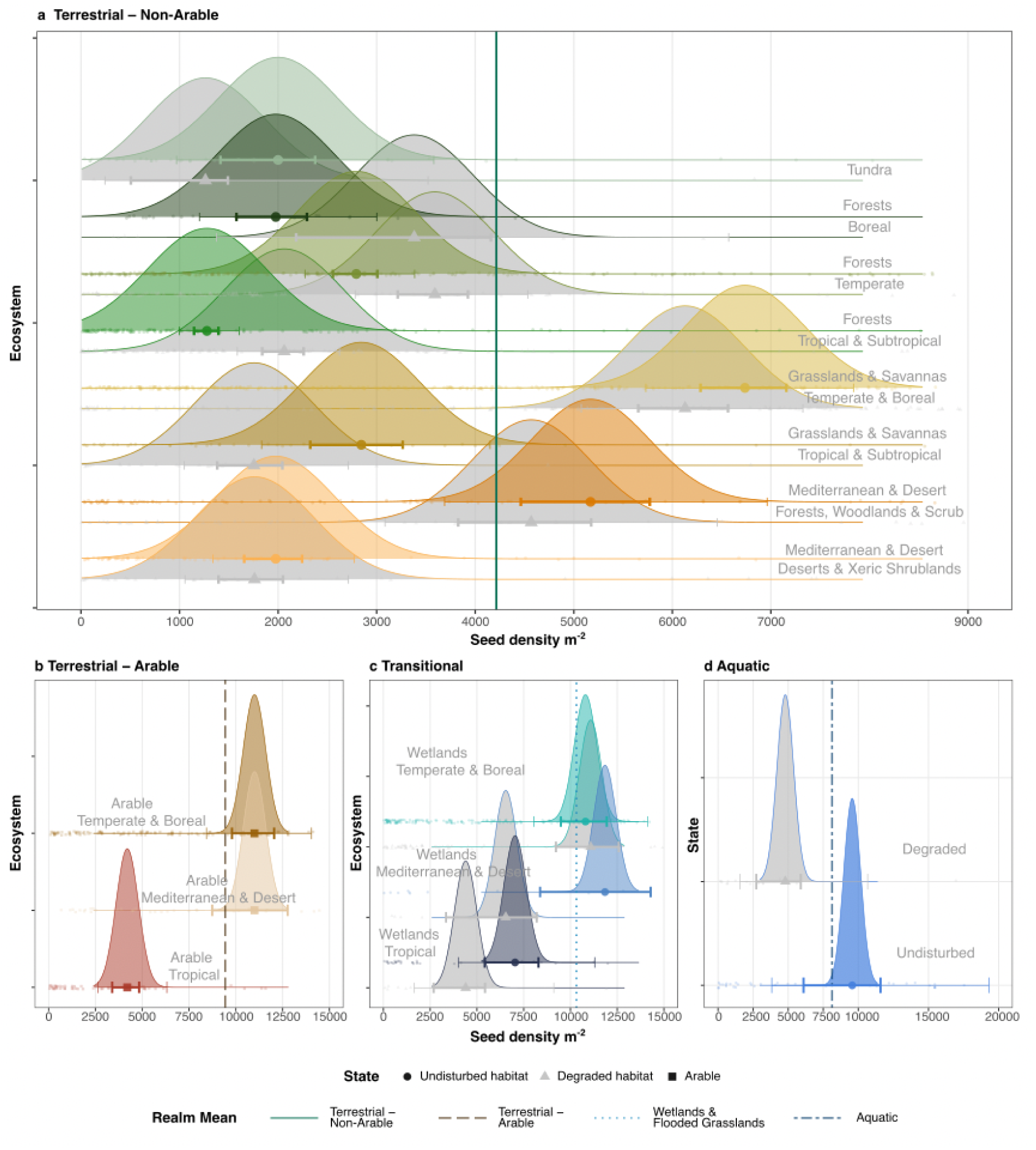
Density of seeds m^−2^ in the soil seed bank across ecosystems. Panels **a-d** represent realms, with the terrestrial realm separated into arable and non-arable systems. Each ecosystem is represented by two density curves and point-and-whiskers, showing soil seed bank density in undisturbed (coloured curves and circular points) and degraded (grey curves and triangular points) ecosystems. Note that arable systems are not separated into disturbed and undisturbed and have only two curves per ecosystem and a square point. Large points show the mean densities, surrounded by 50% (thick) and 90% (thin) credible intervals. Density curves represent 24 000 predictive draws of the posterior distributions of seed density, within each ecosystem and degradation group. Background jittered points represent empirical estimates of seeds density within sampled soil banks. Thick vertical lines in each panel represent the mean fixed effect value across **a** Terrestrial (non-arable records), **b** Terrestrial (arable records), **c** Transitional and **d** Aquatic realms.

### The effect of habitat degradation

In all ecosystems, habitat degradation always resulted in a lower estimated species richness at the 15 m^2^ scale in terms of predicted values, but again, despite this consistency, the difference was only clear in mediterranean forests, woodlands and scrub according to the 50% credible intervals. At the 0.01 m^2^ scale, the predicted values were often the same in undisturbed and degraded ecosystems. Compared to species richness, habitat degradation had much more variable effects on seed density m^−2^ (Figure 3, Figure 4). Broadly speaking, degradation was more likely to result in lower predicted seed densities compared to undisturbed habitats in open ecosystems, with clear differences (50% CIs) for tropical grasslands, mediterranean and desert wetlands and aquatic ecosystems. On the other hand, seed density estimates were generally higher with degradation in forest ecosystems, with clear differences in both temperate and tropical forests (50% CIs). No ecosystems exhibited clear differences in richness or density with degradation according to the 90% CIs.

**Figure 4:**
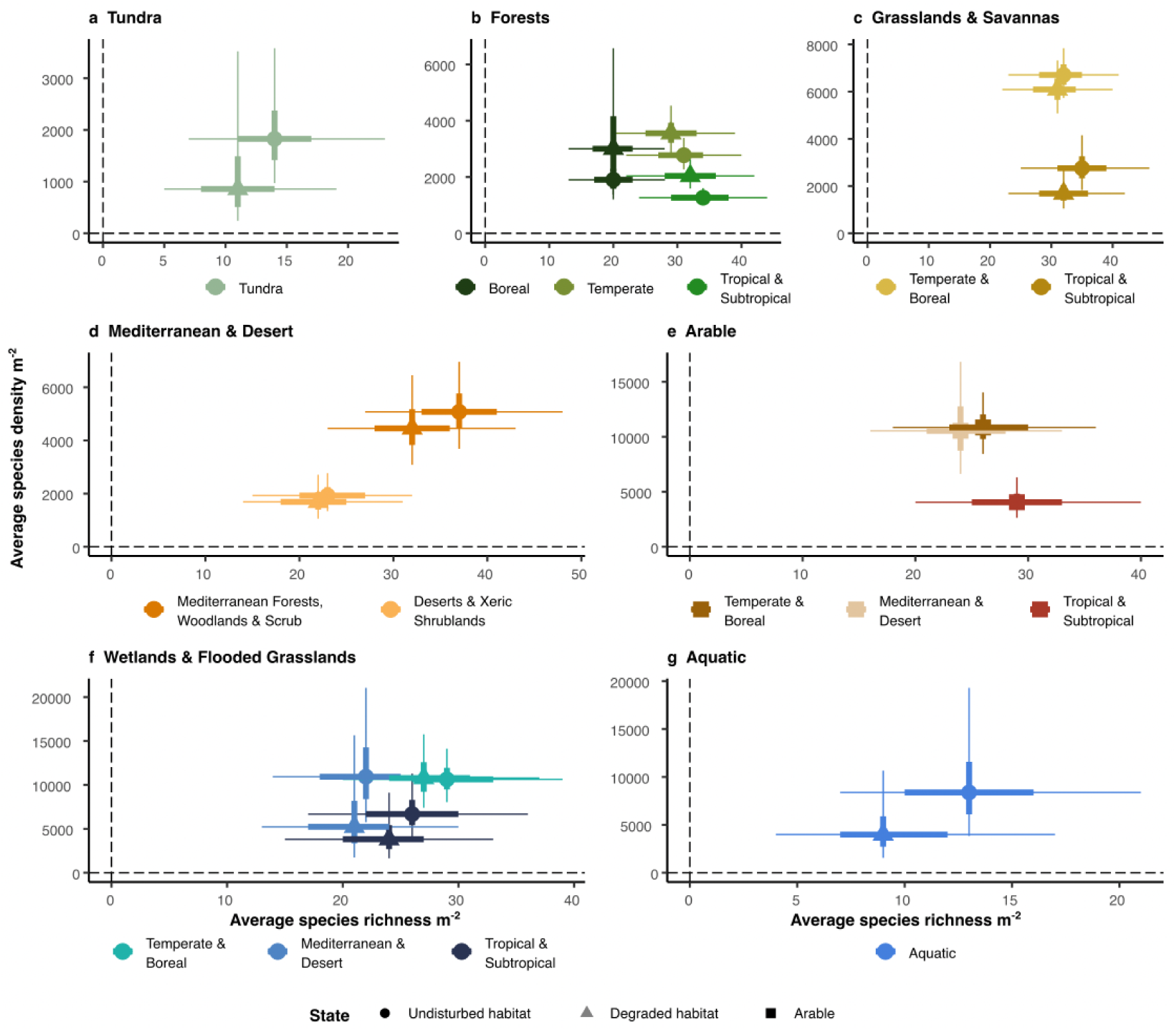
The joint relationship between predicted species richness m^−2^ and seed density m^−2^ in the soil seed bank. Panels **a-g** represent biomes. Each ecosystem is represented by two sets of and point-and-whiskers. Points show the mean estimated species richness and seed density m^−2^, surrounded by 50% (thick) and 90% (thin) credible intervals. Circular points denote undisturbed and triangles denote degraded ecosystems. Arable systems are not separated into disturbed and undisturbed and have one (square) point with whiskers per ecosystem.

### Divergent patterns of richness and density

In many cases, relative patterns of richness and density in the soil seed bank differed both within and among ecosystems, and in response to degradation. Despite relatively high species richness, seed bank density was generally low in tropical systems. Tropical forests had the lowest density across undisturbed ecosystems, being lower than all eleven other undisturbed ecosystems (no overlap of 50% CIs, of which eight had no overlap of 90% CIs). Undisturbed tropical wetlands, forests and grasslands had a lower density than their equivalent ecosystems in temperate and boreal regions (50% CI). Disturbed tropical grasslands were similarly low in seed density, while arable ecosystems in tropical regions were lower than the equivalent systems in other regions (90% CI). Otherwise, arable systems combined relatively low richness with high seed density. Mediterranean and desert, and temperate and boreal arable ecosystems were predicted to have more seeds in the soil than all other ecosystems in the terrestrial realm (50% CIs), while arable ecosystems often had lower richness at the gamma scale than other degraded ecosystems within similar ecoregions (for example, temperate and boreal arable ecosystems had lower richness than temperate forests and temperate and boreal wetlands; 50% CIs). On the other hand, mediterranean forests, woodlands and scrub combined high species richness with relatively high seed density (higher than all other forest ecosystems, 90% CI), while tundra combined its low species richness with low density, especially when degraded, where density was lower than all other degraded ecosystems (50% CI).

## Discussion

Although there was a high uncertainty in our results, we found that across the world’s biomes and ecosystems, patterns in species richness in the seed bank often diverged from patterns of seed density. Strikingly, we also found that the richness of soil seed banks appeared higher in undisturbed compared to degraded habitats, suggesting how across the world’s ecosystems, soil seed banks also reflect the ongoing depletion of biodiversity as a result of anthropogenic activities^23,34^. This indicates that the potential for seed banks to contribute to the recovery and restoration of degraded ecosystems might be limited. Seed density, on the other hand, was not always lower in disturbed habitats, showing how different aspects of biodiversity can show diverging patterns in response to global change pressures^35^. Our estimated global average of more than 5000 seeds m^−2^ also means that in the broadest sense, if you put your finger to the earth, you are likely to be pointing at a seed. The high density of seeds in the soil is a testament to the formation of soil seed banks as an important component of plant population and community dynamics, and a valuable strategy for many plant species to facilitate establishment following disturbance^3,6,36^.

Our analyses revealed substantial variation in the diversity of the soil seed bank among the world’s ecosystems, identifying divergent patterns between species richness and seed density. Although the broad relationship between total abundance and species richness is well-established^35,37^, our understanding of plant biogeography and trait-environment relationships helps us to interpret these results and put them into context. Overall, species richness in the soil seed bank reflected the latitudinal gradient in biodiversity^29,30,38^. However, while tropical and mediterranean ecosystems exhibited relatively high richness compared to temperate, boreal and tundra regions, the former areas differed strongly in terms of seed density. Mediterranean forests and shrublands exhibited both high richness and density in the seed bank (Figures 2-4, Table S3). In regions with warm climates and frequent extreme climatic events, the soil seed bank can play an important role in vegetation recovery and stability of biodiversity^39,40^. Therefore, the high plant biodiversity in these regions^41^, together with the large proportion of species producing persistent seeds^42^, results in the relatively large and rich seed banks in these regions. On the other hand, tropical ecosystems host relatively few species that produce seeds that are able to survive desiccation^13,14^, which is a vital aspect of longevity in the soil. Moreover, the warm and often moist conditions combined with high diversity of other taxa mean that seeds of species that are able to form seed banks are subject to degradation, predation and attacks from pathogens^43–45^. Together, these mechanisms result in low soil seed bank density, but the high overall diversity in the vegetation means that species richness is still high relative to other ecosystems.

The relatively low richness in the soil seed bank in temperate and boreal regions matched expectations relating to broad gradients in biodiversity, and previous work suggests that this effect may be compounded by lower proportions of species in these regions producing persistent seeds compared to lower-latitude species^12^. In these regions, the divergence in patterns of richness and density was more apparent when comparing ecosystems, with density clearly higher in grasslands than forests. The vegetation in the temperate and boreal zones can be considered more stable compared to other regions, and this is especially true in forests. Because seed banking is expected to provide a competitive advantage in more dynamic systems^46,47^, it follows that these relatively stable systems are instead dominated by long-lived perennial species, for which regeneration via the soil seed bank is a less important strategy than for example clonality^48^. In alpine and tundra environments, low seed bank density and a high proportion of clonal species could also be expected, in this case due to a low diversity and density of adult individuals, combined with the risks of seed production during short growing seasons^49,50^.

Our results reveal an imprint of anthropogenic activity on soil seed banks worldwide, and further variation in the relationship between richness and density. While richness was consistently lower in degraded systems, reflecting patterns in degraded habitats in the above-ground vegetation^51–53^, the effect on density was more variable. Unsurprisingly, arable soils contained among the highest number of seeds per square metre (Figures 3 & 4). The high magnitude and frequency of disturbance in arable fields supports relatively species-poor communities of opportunistic ruderal “weed” species whose traits include both high seed longevity in the soil and high seed production^20,21^, and that can be problematic for agricultural production^54–56^. In forest ecosystems, lower richness in the seed bank was often accompanied by higher seed densities. Again, this can be related to plant species’ adaptations to disturbance. While undisturbed forests are relatively stable systems, degraded forests often retain the imprint of the more dynamic management regimes that they are experiencing or recovering from, and therefore their seed banks reflect plant communities that are adapted to such highly-disturbed conditions^57–59^. However, in grasslands and other habitats, degradation generally appeared to result in both reduced richness and density in the soil seed bank. Our findings are concerning, matching general trends of reduced biodiversity in the established vegetation over time^60,61^ (but see ref ^62^), and have consequences for the potential of passive restoration to reverse biodiversity losses^63,64^.

In addition to detecting differences in species richness in the seed bank among ecosystems, we also found differences in the accumulation of richness in space. Our findings support those of Vandvik et al.^25^, who observed that species accumulate more rapidly in alpine and subalpine grasslands than in boreal zones (empirical), and in forests compared to grassland and heathland (literature), and we extend the results to include other global biomes. We found that richness estimates across scales (in our case 0.01 m^2^ to 15m^2^) appeared to show relatively large differences in mediterranean and tropical ecosystems compared to temperate ecosystems, demonstrating how species’ spatial distributions affect how we interpret relative differences in biodiversity across regions (Table S3). These findings have implications for how we understand biodiversity in soil seed banks at both local and regional-global scales.

At local scales, we found that differences in richness across ecosystems, but critically also between undisturbed and degraded ecosystems, were marginal to non-existent at the 0.01 m^2^ scale. This shows that making comparisons at small spatial grains could inhibit our ability to detect important differences in biodiversity that might be relevant for conservation management. At larger spatial scales, comparisons among regions can depend on the spatial grain of predicted richness. This has previously been shown in the aboveground vegetation, where Sabatini et al.^30^ identified a clearer gradient in species richness at large spatial grains, while Keil & Chase^29^ found that areas of relatively high beta diversity in trees – where richness was relatively low at smaller scales and relatively high at larger scales – to be concentrated in low latitudes. Our modelled patterns of soil seed bank richness at the 0.01 m^2^ scale were broadly similar to those found by Yang et al.^26^, although their findings of low richness in the tropics and (predicted) higher richness in northern regions were not supported when considering larger spatial scales.

The large number of soil seed bank studies that have been carried out across the world has allowed our synthesis to cover a greater global extent than many large reviews of biodiversity, which often suffer from biases relating to the absence or restricted availability of data from some regions^65–67^. However, our study may still suffer from other biases, which we discuss here. First, seed banks have not been sampled randomly across the world, but in places where scientists have been interested in finding and identifying seeds in the seed bank^68^. This means that our results rest on the assumption that seed banks were not consistently sampled in areas that are relatively rich or poor in seeds or species, compared to areas that have not been sampled. Second, the database is made up of studies using different sampling efforts to sample the soil seed bank. To address variation in sampled area, we used an approach (ref ^29^ ,Methods), whereby the putative drivers of variation in biodiversity are allowed to vary with sample area in our statistical models, meaning that not only the direct effect of area is controlled for, but also the fact that species-area relationships can also vary across ecosystems and in response to degradation^28,69^. While we also controlled for sampling extent (number of sites) in our models, around half of all observations in the database were represented by a single site. It is well-known that sampling multiple sites per treatment gives improved confidence in ecological findings (sample replication^70^), and we echo previous calls for seed bank sampling to improve representativeness^71,72^. Third, there was a lack of balance in the number of observations across ecosystems, with some better represented than others. Although we hierarchically modelled each biome separately, the distribution of observations among ecoregions and between undisturbed and degraded observations were often uneven. This will have affected our statistical power, increasing uncertainty in our model estimates, particularly where sample sizes were low.

There is another aspect of the study that will have contributed to the uncertainty in our results that is not related to the initial data collection. Our binary categorisation of ecosystem degradation represents a major simplification of the effects of degradation on biodiversity. While we are convinced that such simple classifications can be useful and informative in large-scale studies (see e.g. refs ^23,63^), our general lack of clear differences between undisturbed and degraded ecosystems are likely related to the fact that systems undergoing environmental change are biased towards increasing richness over time^73^. That is, recently degraded or restored ecosystems are likely to contain both colonising species representing the new regime, and species representing the previous regime, which have not yet disappeared. This is especially relevant in the context of seed banks, which exist to buffer short-term environmental heterogeneity, and whose communities often reflect previous habitat conditions^74,75^. In more long-term degraded ecosystems, we may expect the negative effect on seed bank richness to be more clear. It is also important to note that the seed bank dynamics of – for example – a young post-agricultural forest are likely to be quite different from those of a mature plantation, both of which would be classed as degraded forests here. Therefore, while our results may be broadly representative of average effects of degradation at the global scale, the understanding of specific systems undergoing degradation or restoration will require more detailed investigation.

What can we learn from studying the soil seed bank? In addition to the continual publication of studies investigating soil seed banks at local scales, there has been an upswing in the synthesis of large datasets to understand broad spatial or ecological patterns relating to soil seed banks. This includes the macro-environmental determinants of small-scale richness and density^26^, and the evolutionary history of seed bank persistence^76^. Here, we complement such findings with an understanding of the broad biogeographical patterns in soil seed bank richness and density, as well as the spatial scaling of species richness in the seed bank.

Other large-scale work considers the important role that soil seed banks may play in species and community responses to global changes, related to the trade-off related to seeds being able to persist or disperse in space^77^, or the role of soil seed banks in facilitating biological invasions^78^. We believe that the future of soil seed bank synthesis lies beyond simple measures of community richness and density, or categorisation of species as having or not having persistent seeds. Collating a large number of datasets including community data from the soil seed bank and established vegetation – not forgetting to standardise according to spatial scale – across biomes and systems (e.g. ref ^24^), would allow us to delve deeper into a mechanistic understanding of the role of soil seed banks, biodiversity, restoration and responses to global change.

## Methods

### The global seed bank database

To study the effects of spatial scale, habitat type and quality on soil seed banks, we used a global database of seed bank richness, abundance and density^31^. Briefly, the database is based on a Web of Science search using the search term “seed bank*” OR seedbank*, including papers included in the Web of Science until the end of 2019 (a few papers returned by the search were subsequently published in journals in 2020). All resulting papers were checked for being both community seed bank studies (i.e., taking soil samples and counting and identifying all seeds, rather than focussing on one or more specific species), and containing information about species richness (number of species), seed abundance (number of seeds), and/or seed density (number of seeds per unit area or volume) in the seed bank. Most of the papers were in English, but data were also extracted from papers published in, for example, Spanish and Portuguese (often South American studies), as well as French and German. Due to a large data gap, the database was supplemented by an additional database and literature search featuring studies published in Russian and carried out in a number of former Soviet states. In total, this resulted in 7224 unique studies to be checked, and 1442 studies from which records could be extracted or obtained. A <u>record</u> refers to a row in the database, and represents a known number of soil samples with specified (by the study authors) geographical coordinates. Depending on the detail in which study location and design, and seed bank results were reported in a specific study, a record can represent one or more locations or environments. Each record can also provide one or more commonly recorded seed bank metrics (mainly total species, total number of seeds, seeds m^−2^ density), which we term <u>observations</u>. A study can provide multiple observations by reporting more than one metric, and/or by (for example) reporting metrics across several distinct records (i.e. different locations or environments). In total, the 1442 studies provided 8087 observations across 3096 records (Figure 1, Table 1, S1). The database includes soil seed bank data from 94 countries and all seven continents. Full details regarding which data were extracted and how are available in Auffret et al.^31^. Below, we describe only the variables in the database that were used for this analysis.

### Summary information and data variation in the global seed bank

From the 1185 studies and 2585 observations contributing values of seed density, the total number of seeds or seedlings counted across all studies was 35 932 626. Five studies (or 0.2% of all observations) found no seeds within samples. This included an investigation of aquatic systems in Canada^79^, deserts and xeric shrublands in Egypt^80^, temperate forests in Germany and the Netherlands^81,82^, and tundra in Antarctica^83^. Ten studies observed only one species in their seed bank samples, while the maximum was 246 species in 88 m^2^ of soil across 156 arable sites in the midwest of the United States^84^. The minimum (non-zero) observed density of seeds in the seed bank was 3 seeds m^−2^ in a tropical forest in South Africa^85^, with the maximum being 501 119 seeds m^−2^ observed in a study from temperate forests in Germany^86^. The minimum number of seeds greater than zero recorded in a soil seed bank observation was two seeds in two studies in temperate forests in the United States, and tropical and subtropical forests in Indonesia with 0.67 seeds m^−2^ and 0.16 m^−2^ of soil sampled, respectively^87,88^, with the maximum being 5 907 132 seeds in samples taken in arable systems in 462 m^2^ soil from 924 samples across 22 sites in the United Kingdom^89^.

Naturally, such raw maximum and minimum values are at least partly an outcome of each investigation’s scientific focus and study design. Indeed, there was a large variation in sampling effort across records. The minimum total number of samples was one sample, used in 19 studies, with the maximum being 114 000 samples taken in arable systems representing 224 m^2^ of soil sampled across 228 sites in China^90^. The minimum total sampled area was 0.0015 m^2^ (5 sites, 15 total samples) in temperate broadleaf and mixed forests from the United States^91^, with the maximum being 816 m^2^ (20 sites, 20 400 samples) in another large arable study in the UK^92^. The minimum number of sites was one, representing the majority of our samples (1832 observations from 692 studies), with the maximum being 595 sites (total sampled area of 59.5 m^2^) in arable fields across Finland^93^.

### Modelled variables

From the global seed bank database, we extracted the two most common metrics from the soil seed bank literature as response variables: [1] the total number of species recorded in a reported area soil seed bank sampled; [2] the density of seeds m^−2^ in the soil. Soil seed bank studies often involve the collection of multiple samples of soil with a soil corer. These samples are then usually pooled, in order to account for the spatial patchiness of species and seeds in the soil^72,94,95^. Samples can also be collected (and pooled) at several sites in order to capture variation across different locations representing the same habitat or treatment category, often described as ‘replicates’ within a study. We assume that authors of component papers chose a combination of soil samples and sites that they expected would characterise the soil seed bank in their study system. In our analysis, we used the total sampled area in m^2^ or the sample grain as our measure of sampling scale, calculated as the size of the individual soil samples multiplied by the total number of samples taken for a particular observation. We also included the number of sites contributing to each observation as a measure of geographical extent. While this is an imperfect measure, we found it to be a parsimonious covariate, as well as the only such metric that was available in all records. More than half of our observations represent samples collected within a single site (n = 4763).

In order to categorise the global seed bank observations to an ecosystem in a systematic way, we used a combination of authors self-declared habitat descriptions coming from the global seed bank database (aquatic, open grassy systems, forest, arable, or wetland), broad categories of the WWF terrestrial ecoregions based on an observation’s geographic coordinates^32^, as well as realms in the fashion of the function-based typology for the Earth’s ecosystems^33^. Observations with habitat defined as arable or aquatic were placed in an arable and aquatic ‘biome’, respectively. Habitat and biome were used to categorise all observations into one of 15 categories (Table 1), with the aim to maintain ecologically-relevant distinctions of ecosystems (the combination of biome and ecoregion) while also including a reasonable number of observations within each group. Each ecosystem was then assigned into either the aquatic, terrestrial or transitional realm. Within the terrestrial realm, we assigned biomes as tundra, forests, grasslands, mediterranean and desert, and arable systems, while we included all aquatic observations within an aquatic biome within the aquatic realm. The transitional realm included wetlands in the mediterranean and desert, temperate and boreal, and tropical ecoregions (Table 1). Finally, the database labelled non-arable observations as coming from undisturbed or degraded habitats. Habitat degradation was included as a binary classification within each ecosystem, based on descriptions from the component studies. During compilation of the database, the contemporary habitat being assigned to one of the habitat categories listed above, with an additional ‘target habitat’ was also added where necessary. Target habitat refers to the ideal natural or semi-natural habitat of the system, according to the study’s authors, interpreted from site descriptions and/or due to the common practice in the seed bank literature of comparing assemblages across habitat gradients. If there was no entry for target habitat for a record in the database, we used the (contemporary) habitat entry and the record was labelled as being undisturbed. When there was an entry for target habitat, this was used for the habitat definition and the record was labelled as degraded. Note that the target habitat can have the same category as the contemporary habitat if it is still – for example – a forest, but in a degraded state. Examples of degraded habitats include drained wetlands, early-mid successional secondary forests, abandoned semi-natural grassland or any habitat under restoration, see Auffret et al.^31^ for further details.

### Statistical Models

For each response (i.e. species richness and seed density) and biome (e.g. terrestrial grasslands and transitional wetlands) we fit a univariate mixed effects linear model. This amounted to 14 models in total (two responses × seven biomes; Table S2), allowing estimates to vary independently for each biome without well-sampled biomes and realms (e.g. temperate forests) influencing the estimates of undersampled biomes and realms (e.g. tundra, aquatic). In general, if there were different ecoregions represented within a biome then these were treated as categorical fixed effects that interacted with habitat degradation. If not (e.g. tundra), then degradation was the only categorical fixed effect. In the arable biome, there was no fixed effect for degradation, because we consider arable to already represent a degraded system. For each model, intercepts were allowed to vary for every record, nested within study identity as random effects.

In the species richness models, we quantified richness as a function of sampling area across the different biomes by fitting total sampled area (m^2^; sample grain) as a continuous predictor, ecoregion as an interacting categorical predictor (where relevant), degradation as an interacting categorical predictor (a three-way interaction where relevant), and the number of sites as a continuous covariate as a proxy for study extent. This approach, allowing drivers of diversity to vary with sampling grain within a single model, has been found to effectively unify disparate results from previous studies, enabling efficient integration of heterogeneous biodiversity data^29^. We assumed a Poisson distribution with a log-link function and we log-transformed and centred the total sampling area and the number of sites. We set the lower bound of slopes to be zero as informative prior information. This prevented uncertainty estimates for shallow slopes (aquatic realm) from overlapping with negative values, which does not logically hold. Seed density models included only degradation interacting with ecoregion as fixed effects, and assumed a lognormal distribution. For all model specifications and details see Table S2.

For Bayesian inference and estimates of uncertainty, we fit models using the Hamiltonian Monte Carlo (HMC) sampler Stan^96^ using the R package brms (citations for all software are given at the end of the Methods). We fit all models with four chains, differing iterations and differing assumed distributions (Supplementary Information, Model Details Table S2). We used weakly regularising priors, visually inspected HMC chains for convergence, and conducted posterior predictive checks to ensure the model represented the distribution of the data for each (Supplementary Information, Table S2, Figure S1a-g). We selected model structure and distributions based on this and on model performance (Rhats, sample sizes) (Supplementary Information, Table S2 Figure S1a-g).

### Quantifying scale-dependent richness

Relative differences in local species richness (alpha diversity) between habitats or biomes do not always match patterns of richness when larger areas are considered (gamma diversity). This indicates spatial variation in beta diversity (Whittaker’s beta = gamma / alpha)^97^. To investigate this divergence between scales, we first predicted the overall average species richness at two spatial scales for every biome using the species richness models to keep richness estimates for every biome consistent. We predicted richness at 0.01 m^2^ (alpha diversity), and at a larger scale, gamma diversity at 15 m^2^, keeping modelling parameters constant. These are arbitrary values, but we consider 0.01 m^2^ to be a commonly sampled size in soil seed bank studies, that falls within the interquartile range of seed bank sampling in the database^31^ and has been previously used for pattern prediction of soil seed bank richness and density^26^. Gamma diversity at 15 m^2^ was chosen as the largest scale that we felt confident predicting across biomes, given the input data and model outputs. In the main results we report diversity at both alpha and gamma scales assuming each observation was carried out at one site, as this represents the majority of our values, and acts as our baseline estimates for diversity.

We estimated the average number of seeds m^−2^ globally for observations within non-disturbed natural terrestrial, arable, wetland and aquatic realms. To do so, we combined estimates of the fixed effect (i.e. fitted values) values across all models (i.e. biomes) from each realm, and predicted the mean seed density m^−2^ across all fitted values from all models within each broad group. Finally, to look at the joint-relationship between the density of seed in the soil seed bank and species richness, we predicted species richness at the 1 m^2^ scale used for seed density for direct comparison between the two. We predicted richness at this third scale 1 m^2^, following the same methods at, as for our alpha and gamma diversity. For all data cleaning, management, wrangling and analyses, we used the R Project for Statistical Computing version 4.4.0^98^ (R Project). To handle data, quantify metrics, summarise results, and visualise data and estimates we used the brms, tidyverse, bayesplot, and ggplot2 packages^99–102^.

## Data Availability

The global seed bank database is published at https://doi.org/10.5878/bvs7-gk47 and is fully described in ref ^31^.

## Code Availability

The code used for the analysis is available at https://github.com/Global-Seed-Banks/Seed-Banks

## Acknowledgements

We thank the Landscape Ecology unit at SLU, and the Biodiversity Synthesis and Physiological Diversity groups at iDiv for feedback on the project idea, data analysis and presentation. AGA is supported by the Swedish research councils VR (2020- 04276) and Formas (2023-02457). EL was supported by the German Centre for Integrative Biodiversity Research (iDiv) Halle-Jena-Leipzig, funded by the German Research Foundation (FZT 118, 202548816). PK was funded by the European Union (ERC, BEAST, 101044740). Views and opinions expressed are however those of the author(s) only and do not necessarily reflect those of the European Union or the European Research Council Executive Agency. Neither the European Union nor the granting authority can be held responsible for them. VO thanks RSF for financial support (# 25-14-00032). ED was supported by Formas (2023-01507), as were IK (2022-01752, via Biodiversa+ BiodivProtect) and JP (2018-00961 and 2022-01803).

