## Supplementary information for "Divergent patterns of richness and density in the global soil seed bank"

### <u>Contents</u>

**Data**
**Table S1**. Summary of the Global Seed Bank Database

**Model Specifications & Checks**
**Table S2.** Details on each univariate hierarchical mixed effects model for main analysis
**Figure S1**. Predicted values for each richness (a) and density (b) biome model

**Main Model Estimated Results**
**Table S3.** Richness-area slopes.
**Table S4.** Summary of model estimates of seed density
**Table S5:** Predicted estimates of diversity at different scales
**Table S6. Average soil seed bank densities per realm.**

**Supplementary Figures**
**Figure S2.** Species richness in the soil seed bank per biome

### Data

**Table S1. Summary of the Global Seed Bank Database**. Number of studies checked for data, total number of studies with usable data, total number of records, and observations. By ‘observation’, we mean a single value for a metric describing the seed bank community (such as richness, density or abundance, see below). We do not count the ratio within total observations, as we derived it from reported observations. Records are a unique sampled area reported within a study. A study can provide multiple observations by reporting more than one variable, and/or by (for example) reporting variables from several distinct records, which could be locations or habitat types. The number of total observations in the database differs from that used in the statistical models because for species richness and seed abundance, the study needed to also report the area sampled and for seed density we removed zeros. We report more detail on that, and on other model sample sizes further down in the data section and under ‘Model Specifications’.

| Data | n | Number of observations used in each statistical model |
|---|---|---|
| Total number of studies checked from literature | 7224 | |
| Total number of studies with usable data | 1442 | |
| Total number of records | 3096 | |
| Total number of observations (richness, abundance and density combined) | 8087 | |
| Total species (richness) | 2872 | 2792 |
| Total seeds (abundance) | 2585 | 2569 |
| Number of seeds/m2 (density) | 2630 | 2626 |

### Model Specifications & Checks

**Table S2. Details on each univariate hierarchical mixed effects model for main analysis**. Poisson distributions with a log-link function were selected for models where the response was species richness. Lognormal distributions were selected for density and ratio models, with zeros removed (observations where zero seeds were found) as the log of zero is unknown. Bulk Effective Sample Sizes are reported. All models resulted in a Rhat of 1. Within a model short-name group, if the formula is blank, it follows the same formula as the first mentioned (tundra, top row of each model group) formulas are only listed after that diverge from that first listed formula.

| Response | Formula | Biome | Figure | N observations | Distribution | Iterations | Effective Sample Size |
|---|---|---|---|---|---|---|---|
| Species richness | richness ~ centred log area * degraded * biome subtype + centred log number of sites + ( 1 \| study / record) | Tundra | 2, S1, S2 | 43 | Poisson | 2000 | 1202-2507 |
| | | Forest | | 734 | Poisson | 2000 | 1399-4424 |
| | | Grasslands | | 882 | Poisson | 2000 | 577-2240 |
| | | Mediterranean & Deserts | | 330 | Poisson | 8000 | 8895-24316 |
| | richness ~ centred log area * biome subtype + centred log number of sites + ( 1 \| study / record) | Arable | | 257 | Poisson | 6000 | 9156-19042 |

| Response | Formula | Biome | Figure | N observations | Distribution | Iterations | Effective Sample Size |
|---|---|---|---|---|---|---|---|
| | | Wetlands | | 472 | Poisson | 2000 | 1406-4129 |
| | richness ~ centred log area * degraded + centred log number of sites + ( 1 \| study / record) | Aquatic | | 73 | Poisson | 2000 | 942-2836 |
| Seed density | seed density m2 ~ degraded * biome subtype + ( 1 \| study / record) | Tundra | 3, 2b | 41 | Lognormal | 10000 | 17277-26499 |
| | | Forest | | 749 | Lognormal | 10000 | 7736-8703 |
| | | Grasslands | | 796 | Lognormal | 10000 | 7024-18898 |
| | | Mediterranean & Deserts | | 300 | Lognormal | 10000 | 13365-18766 |
| | seed density m2 ~ biome subtype + ( 1 \| study / record) | Arable | | 256 | Lognormal | 10000 | 14238-17036 |
| | | Wetlands | | 420 | Lognormal | 10000 | 9620-10391 |
| | seed density m2 ~ degraded + ( 1 \| study / record) | Aquatic | | 63 | Lognormal | 10000 | 10520-18130 |

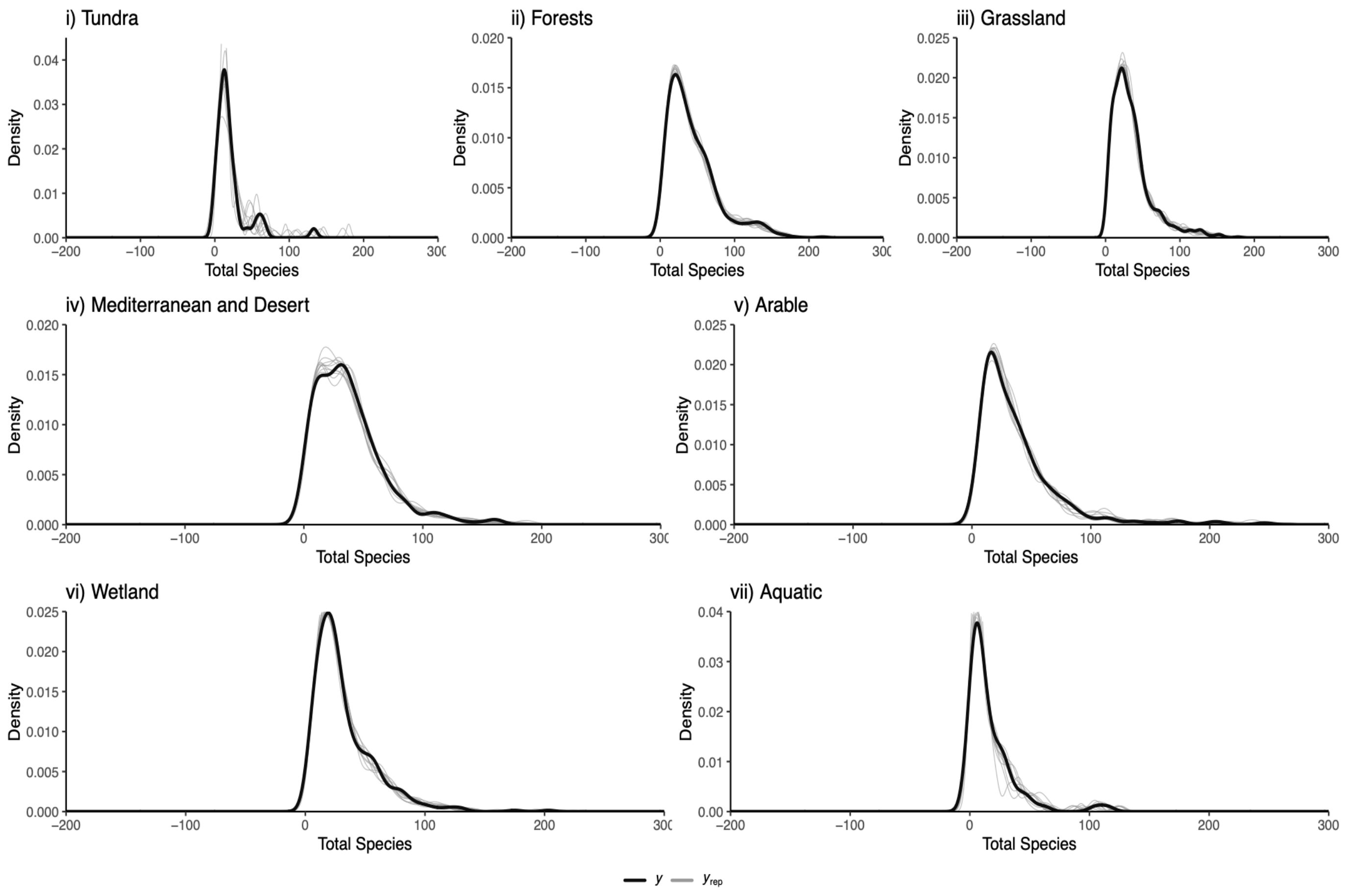


**Figure S1a i)-vii) :** Predicted values for each species richness biome model indicated by the grey lines and observed values of the data indicated by the black line.

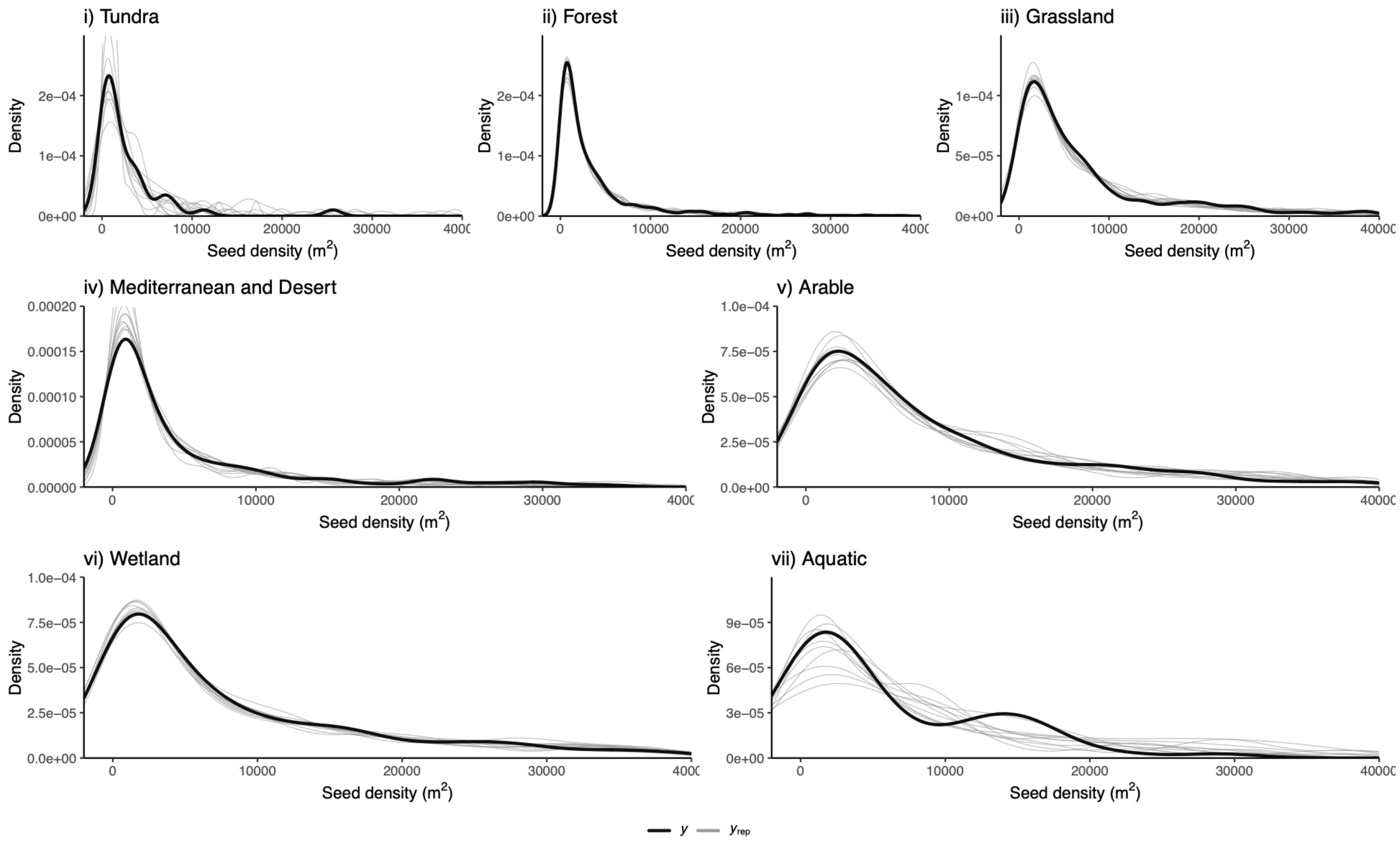


**Figure S1b i)-vii):** Predicted values for each seed density biome model indicated by the grey lines and observed values of the data indicated by the black line.

### Main Model Estimated Results

**Table S3. Richness-area slopes.** Summary of model estimates regarding species richness slope as a function of area. In each column, the model estimate is followed by the lower credible interval and upper 50% credible interval, followed by 90% in brackets. Biomes and Realms presented in the same order as in Figures.

| Realm | Biome | Ecoregion | Degraded | Slope and Intervals |
|---|---|---|---|---|
| Terrestrial | Tundra | Tundra | 1 | 0.1063 (0.0405-0.1519 , 0.0073-0.266) |
| Terrestrial | Tundra | Tundra | 0 | 0.2053 (0.135-0.265 , 0.0649-0.3773) |
| Terrestrial | Forest | Boreal | 1 | 0.1018 (0.0803-0.1261 , 0.0384-0.152) |
| Terrestrial | Forest | Boreal | 0 | 0.1163 (0.0936-0.1412 , 0.0507-0.172) |
| Terrestrial | Forest | Temperate | 1 | 0.1494 (0.1328-0.1657 , 0.1077-0.1925) |
| Terrestrial | Forest | Temperate | 0 | 0.1164 (0.0939-0.141 , 0.0517-0.1682) |
| Terrestrial | Forest | Tropical | 1 | 0.1472 (0.1304-0.1636 , 0.1063-0.1873) |
| Terrestrial | Forest | Tropical | 0 | 0.1136 (0.0919-0.1381 , 0.0491-0.1659) |
| Terrestrial | Grassland | Temperate and Boreal | 1 | 0.0858 (0.0713-0.1004 , 0.0511-0.1202) |
| Terrestrial | Grassland | Temperate and Boreal | 0 | 0.1215 (0.0987-0.1427 , 0.0733-0.1762) |
| Terrestrial | Grassland | Tropical | 1 | 0.0997 (0.0843-0.1153 , 0.0616-0.138) |
| Terrestrial | Grassland | Tropical | 0 | 0.1033 (0.0859-0.1204 , 0.0626-0.1448) |
| Terrestrial | Mediterranean and Desert | Deserts and Xeric Shrublands | 1 | 0.148 (0.1272-0.1697 , 0.0942-0.1994) |
| Terrestrial | Mediterranean and Desert | Deserts and Xeric Shrublands | 0 | 0.2129 (0.1708-0.2497 , 0.1269-0.3187) |

| | | | | |
|---|---|---|---|---|
| Terrestrial | Mediterranean and Desert | Mediterranean Forests, Woodlands and Scrub | 1 | 0.1667 (0.1455-0.1881 , 0.114-0.2191) |
| Terrestrial | Mediterranean and Desert | Mediterranean Forests, Woodlands and Scrub | 0 | 0.1752 (0.1514-0.199 , 0.1152-0.2353) |
| Terrestrial | Arable | Mediterranean and Desert | 1 | 0.0476 (0.0243-0.0672 , 0.0057-0.1008) |
| Terrestrial | Arable | Temperate and Boreal | 1 | 0.0852 (0.0638-0.1054 , 0.0371-0.1357) |
| Terrestrial | Arable | Tropical | 1 | 0.0865 (0.0572-0.1119 , 0.0268-0.1571) |
| Transitional | Wetland | Mediterranean and Desert | 1 | 0.0766 (0.0443-0.1057 , 0.0117-0.149) |
| Transitional | Wetland | Mediterranean and Desert | 0 | 0.1104 (0.0723-0.1449 , 0.0322-0.1998) |
| Transitional | Wetland | Temperate and Boreal | 1 | 0.156 (0.133-0.1791 , 0.1-0.2115) |
| Transitional | Wetland | Temperate and Boreal | 0 | 0.0936 (0.0603-0.1237 , 0.0245-0.1698) |
| Transitional | Wetland | Tropical | 1 | 0.1574 (0.1177-0.1941 , 0.0683-0.252) |
| Transitional | Wetland | Tropical | 0 | 0.1247 (0.0855-0.1595 , 0.0428-0.2206) |
| Aquatic | Aquatic | Aquatic | 1 | 0.0801 (0.0298-0.1151 , 0.0062-0.2026) |
| Aquatic | Aquatic | Aquatic | 0 | 0.131 (0.0753-0.1744 , 0.0329-0.2669) |

**Table S4. Summary of model estimates of seed density$^{-2}$.** In each column, the model estimate (mean) is followed by the lower credible interval and upper 50% credible interval, followed by 90% in brackets. Biomes and Realms presented in the same order as in Figures.

| Realm | Biome | Ecoregion | Degraded | Density and Intervals |
|---|---|---|---|---|
| Terrestrial | Tundra | Tundra | 1 | 856 (506-1491 , 242-3521) |
| Terrestrial | Tundra | Tundra | 0 | 1826 (1415-2376 , 969-3580) |
| Terrestrial | Forest | Boreal | 1 | 1899 (1577-2292 , 1202-3001) |
| Terrestrial | Forest | Boreal | 0 | 3001 (2183-4159 , 1374-6572) |
| Terrestrial | Forest | Temperate | 1 | 2773 (2556-3007 , 2272-3385) |
| Terrestrial | Forest | Temperate | 0 | 3554 (3214-3926 , 2782-4533) |
| Terrestrial | Forest | Tropical | 1 | 1264 (1148-1395 , 999-1604) |
| Terrestrial | Forest | Tropical | 0 | 2036 (1838-2259 , 1586-2622) |
| Terrestrial | Grassland | Temperate and Boreal | 1 | 6709 (6286-7156 , 5731-7838) |
| Terrestrial | Grassland | Temperate and Boreal | 0 | 6090 (5654-6564 , 5075-7324) |
| Terrestrial | Grassland | Tropical | 1 | 2757 (2326-3264 , 1833-4151) |
| Terrestrial | Grassland | Tropical | 0 | 1683 (1383-2043 , 1046-2711) |
| Terrestrial | Mediterranean and Desert | Deserts and Xeric Shrublands | 1 | 1930 (1656-2243 , 1339-2774) |
| Terrestrial | Mediterranean and Desert | Deserts and Xeric Shrublands | 0 | 1691 (1395-2049 , 1055-2711) |
| Terrestrial | Mediterranean and Desert | Mediterranean Forests, Woodlands and Scrub | 1 | 5078 (4463-5771 , 3689-6962) |
| Terrestrial | Mediterranean and Desert | Mediterranean Forests, Woodlands and Scrub | 0 | 4454 (3827-5176 , 3085-6455) |
| Terrestrial | Arable | Mediterranean and Desert | 1 | 10547 (8739-12770 , 6636-16829) |

| | | | | |
|---|---|---|---|---|
| Terrestrial | Arable | Temperate and Boreal | 1 | 10854 (9797-12051 , 8436-14047) |
| Terrestrial | Arable | Tropical | 1 | 4049 (3391-4833 , 2633-6317) |
| Transitional | Wetland | Mediterranean and Desert | 1 | 10934 (8377-14288 , 5760-21058) |
| Transitional | Wetland | Mediterranean and Desert | 0 | 5237 (3349-8196 , 1735-15653) |
| Transitional | Wetland | Temperate and Boreal | 1 | 10635 (9487-11942 , 8036-14127) |
| Transitional | Wetland | Temperate and Boreal | 0 | 10780 (9226-12588 , 7404-15767) |
| Transitional | Wetland | Tropical | 1 | 6683 (5422-8289 , 4001-11296) |
| Transitional | Wetland | Tropical | 0 | 3818 (2688-5433 , 1638-9112) |
| Aquatic | Aquatic | Aquatic | 1 | 3996 (2727-5898 , 1556-10664) |
| Aquatic | Aquatic | Aquatic | 0 | 8385 (6092-11581 , 3833-19307) |

### Predicted values

**Table S5. Predicted estimates of diversity at different scales.** alpha = 0.01 $m^2$, gamma = 15 $m^2$. In the main results, we report the predicted alpha and gamma values for one site. Here, the number after each underscore specifies the predictions at each scale for the 3 levels of number of sites. Table sorted in ascending average alpha richness values at 1 site, average gamma richness values at 1 site and alphabetical order of Biomes.

| Realm | Biome | Ecoregion | Degraded | 0.01 | 1 | 15 |
|---|---|---|---|---|---|---|
| Terrestrial | Tundra | Tundra | 0 | 3 (1-4 , 0-7) | 14 (17,11 , 7,23) | 15 (7-19 , 3-37) |
| Terrestrial | Tundra | Tundra | 1 | 3 (2-5 , 0-8) | 11 (14,8 , 4,20) | 8 (4-10 , 1-18) |
| Terrestrial | Forest | Boreal | 0 | 10 (7-12 , 4-16) | 20 (23,17 , 12,28) | 21 (17-25 , 13-31) |
| Terrestrial | Forest | Boreal | 1 | 10 (7-12 , 4-16) | 20 (23,16 , 12,28) | 20 (16-24 , 12-30) |
| Terrestrial | Forest | Temperate | 0 | 11 (9-14 , 6-18) | 30 (34,27 , 22,40) | 40 (35-46 , 27-55) |
| Terrestrial | Forest | Temperate | 1 | 12 (9-14 , 6-19) | 29 (33,25 , 21,39) | 36 (31-41 , 24-49) |
| Terrestrial | Forest | Tropical | 0 | 13 (10-15 , 7-20) | 34 (38,29 , 24,44) | 44 (38-49 , 31-58) |
| Terrestrial | Forest | Tropical | 1 | 13 (10-16 , 7-20) | 32 (36,28 , 22,42) | 39 (33-43 , 27-52) |
| Terrestrial | Grasslands | Temperate and Boreal | 0 | 14 (11-16 , 8-21) | 32 (35,28 , 23,41) | 29 (25-34 , 19-40) |
| Terrestrial | Grasslands | Temperate and Boreal | 1 | 14 (11-17 , 8-21) | 31 (34,27 , 22,40) | 26 (22-31 , 17-37) |
| Terrestrial | Grasslands | Tropical | 0 | 14 (11-16 , 7-21) | 35 (39,31 , 25,46) | 36 (31-41 , 24-50) |
| Terrestrial | Grasslands | Tropical | 1 | 14 (11-17 , 8-21) | 33 (37,28 , 23,43) | 30 (25-34 , 19-41) |
| Terrestrial | Mediterranean and Desert | Deserts and Xeric Shrublands | 0 | 10 (8-13 , 5-17) | 23 (27,20 , 15,32) | 39 (32-44 , 25-54) |
| Terrestrial | Mediterranean and Desert | Deserts and Xeric Shrublands | 1 | 11 (8-13 , 5-18) | 22 (25,18 , 14,31) | 32 (27-37 , 20-45) |
| Terrestrial | Mediterranean and Desert | Mediterranean Forests, Woodlands and Scrub | 0 | 14 (11-17 , 7-22) | 37 (41,33 , 27,48) | 73 (63-83 , 50-99) |
| Terrestrial | Mediterranean and Desert | Mediterranean Forests, Woodlands and Scrub | 1 | 15 (12-18 , 8-23) | 32 (36,28 , 23,43) | 51 (43-58 , 34-70) |
| Terrestrial | Arable | Mediterranean and Desert | 1 | 13 (10-15 , 7-20) | 25 (28,21 , 16,33) | 18 (14-22 , 10-28) |
| Terrestrial | Arable | Temperate and Boreal | 1 | 12 (9-14 , 6-19) | 26 (30,23 , 18,36) | 23 (18-27 , 13-34) |
| Terrestrial | Arable | Tropical | 1 | 13 (10-16 , 6-21) | 29 (33,25 , 20,40) | 25 (20-29 , 15-38) |

| | | | | | | |
|---|---|---|---|---|---|---|
| Transitional | Wetland | Mediterranean and Desert | 0 | 11 (8-14 , 5-18) | 22 (25,18 , 14,31) | 21 (17-25 , 12-33) |
| Transitional | Wetland | Mediterranean and Desert | 1 | 11 (8-14 , 5-19) | 21 (24,17 , 13,30) | 20 (15-24 , 11-31) |
| Transitional | Wetland | Temperate and Boreal | 0 | 11 (8-13 , 5-17) | 30 (33,26 , 21,39) | 42 (34-48 , 26-60) |
| Transitional | Wetland | Temperate and Boreal | 1 | 11 (8-13 , 5-18) | 27 (31,23 , 18,37) | 35 (28-40 , 21-51) |
| Transitional | Wetland | Tropical | 0 | 9 (6-11 , 3-15) | 26 (30,23 , 17,36) | 42 (33-50 , 24-65) |
| Transitional | Wetland | Tropical | 1 | 10 (7-12 , 4-17) | 24 (27,20 , 15,33) | 30 (24-36 , 17-48) |
| Aquatic | Aquatic | Aquatic | 0 | 6 (3-8 , 1-12) | 13 (16,10 , 7,22) | 15 (9-20 , 4-33) |
| Aquatic | Aquatic | Aquatic | 1 | 5 (3-6 , 1-11) | 9 (12,7 , 4,16) | 9 (5-12 , 2-20) |

**Table S6. Average soil seed bank densities per realm.** Realm mean, median and 50% and 90% upper and lower credible intervals based on the pooled fitted values of each model.

| Realm | median | mean | 50% Credible Interval | 90% Credible Interval |
|---|---|---|---|---|
| Natural Terrestrial | 3590 | 4214 | 3814-4565 | 3364-5217 |
| Arable | 11001 | 9434 | 8143-10508 | 6796-12733 |
| Wetlands | 10802 | 10318 | 8699-11626 | 7115-14481 |
| Aquatic | 9562 | 8124 | 5077-9867 | 3146-16700 |

### Supplementary Figures

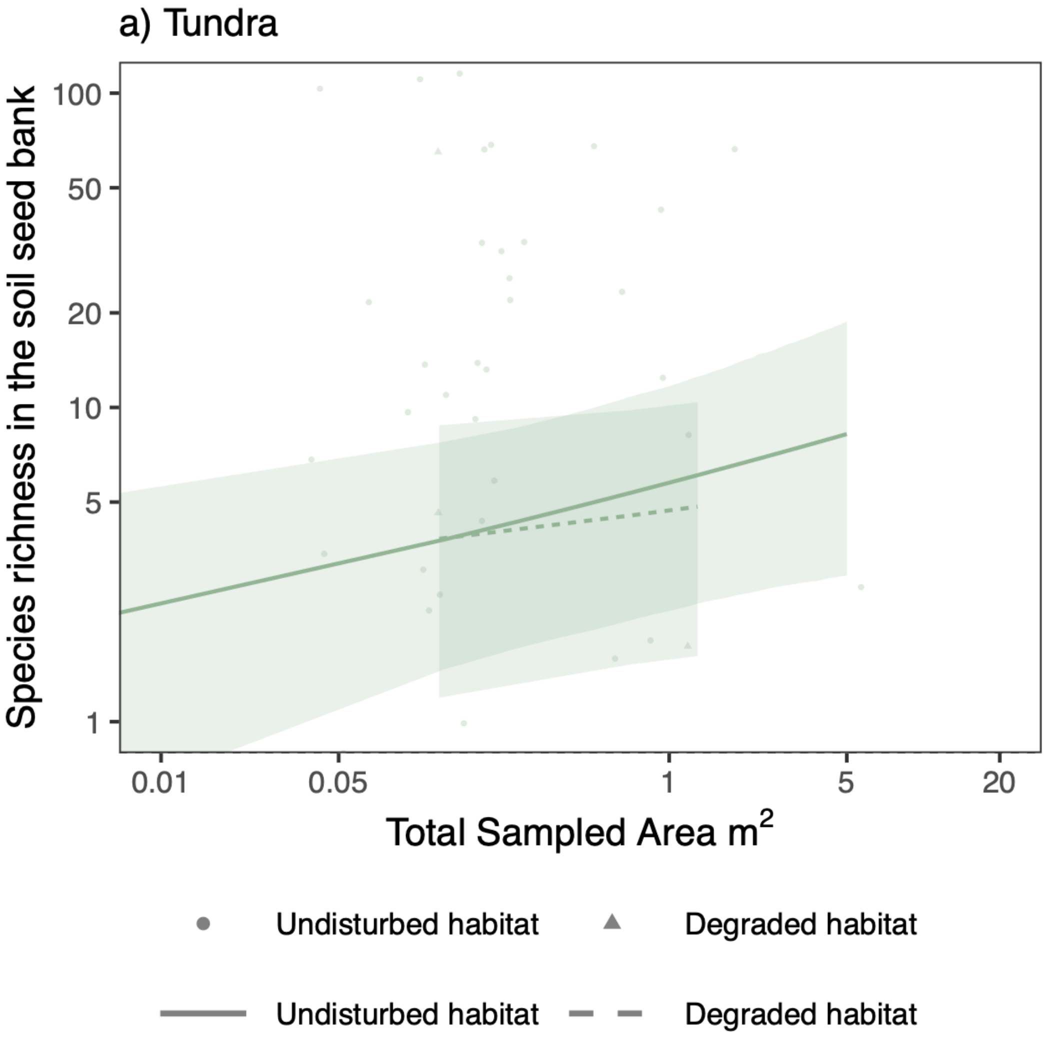


**Figure S2a. Species richness in the soil seed bank as a function of total soil samples ($m^2$)**. Each point and line type represents an undisturbed habitat (circles, solid line), and disturbed habitat (triangles, dotted line). Each point is the species richness reported for a given total area of soil sampled in the literature, each line is the overall estimated relationship between the number of species and sampled area for one site, and the shaded area around lines are the 95% credible intervals.

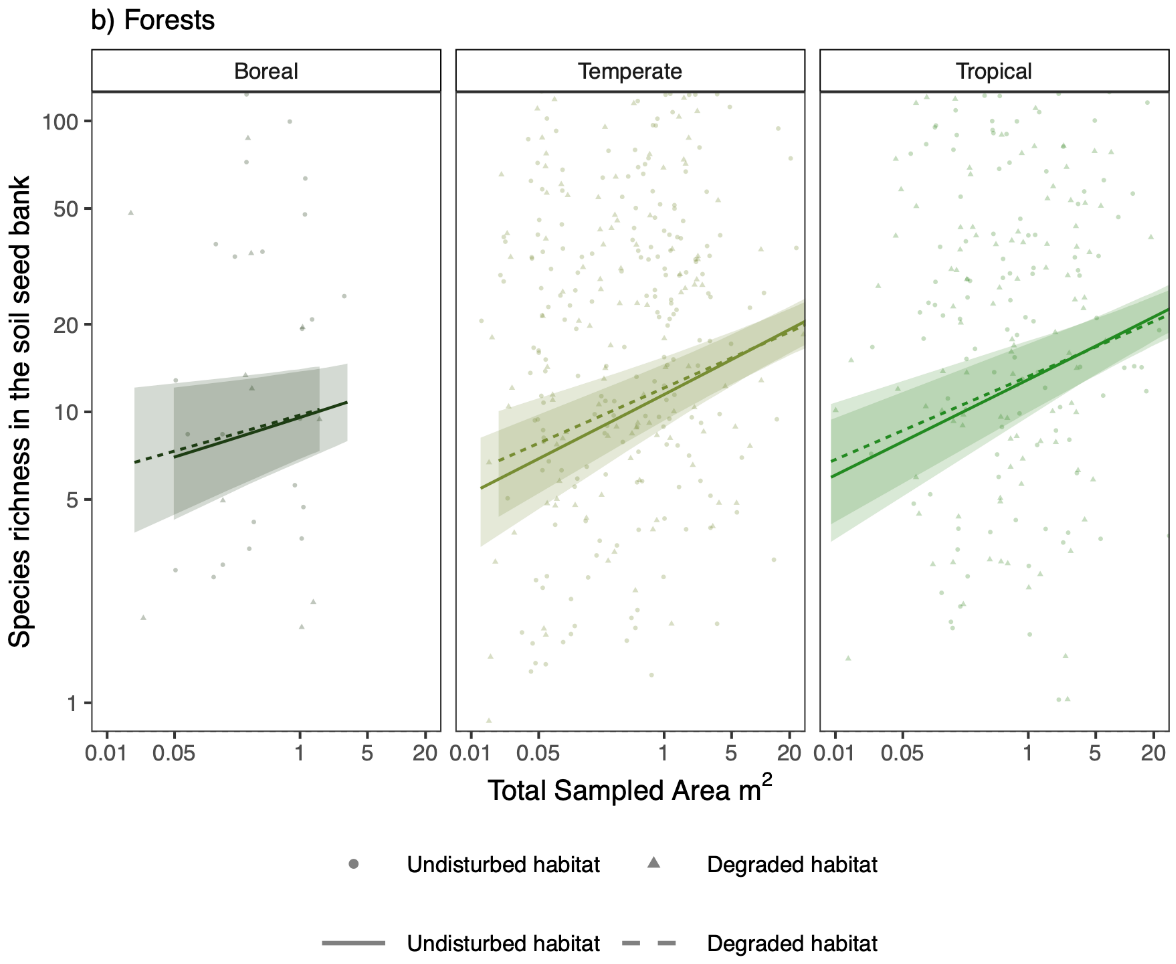


**Figure S2b. Species richness in the soil seed bank as a function of total soil samples ($m^2$).** Each point and line type represents an undisturbed habitat (circles, solid line), and disturbed habitat (triangles, dotted line). Each point is the species richness reported for a given total area of soil sampled in the literature, each line is the overall estimated relationship between the number of species and sampled area for one site, and the shaded area around lines are the 95% credible intervals.

.

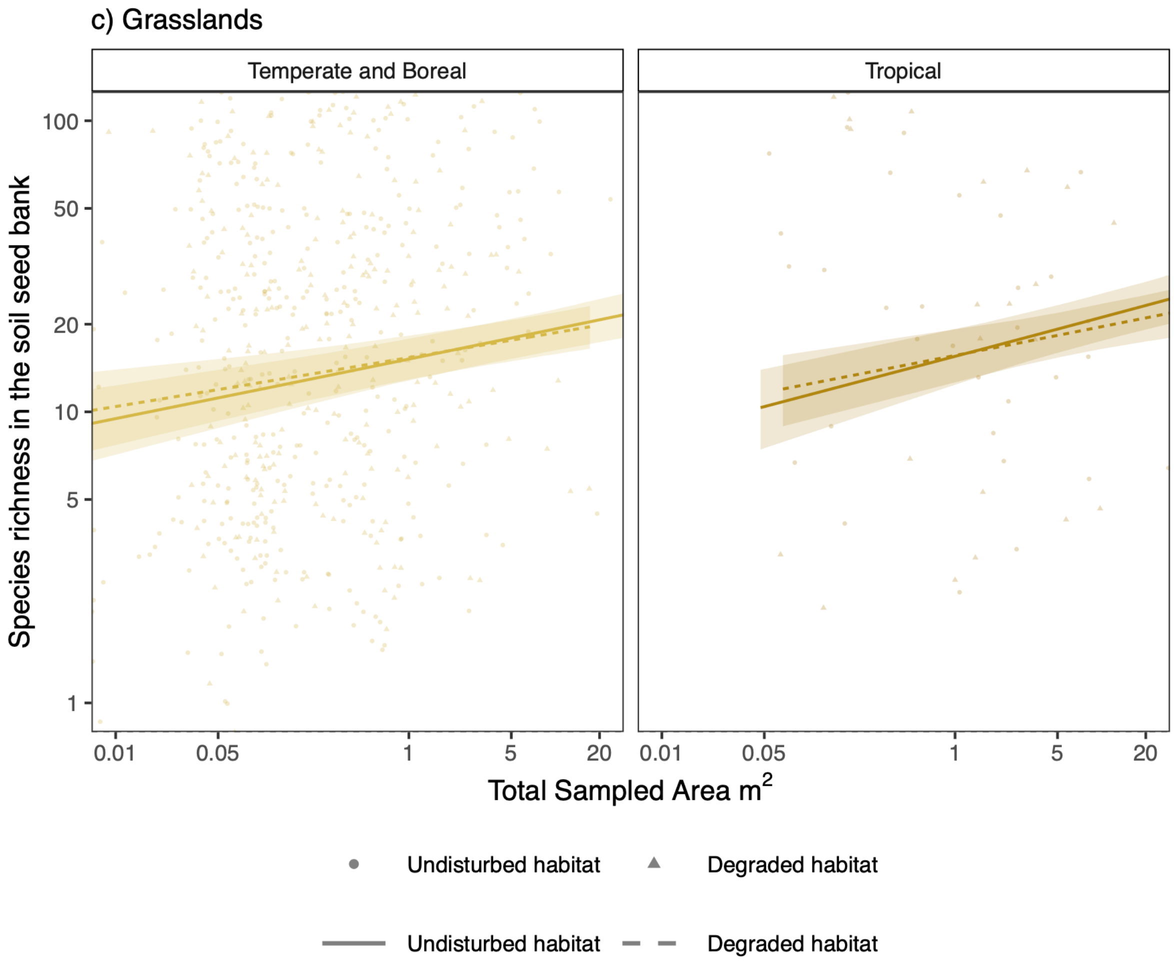


**Figure S2c. Species richness in the soil seed bank as a function of total soil samples ($m^2$).** Each point and line type represents an undisturbed habitat (circles, solid line), and disturbed habitat (triangles, dotted line). Each point is the species richness reported for a given total area of soil sampled in the literature, each line is the overall estimated relationship between the number of species and sampled area for one site, and the shaded area around lines are the 95% credible intervals.

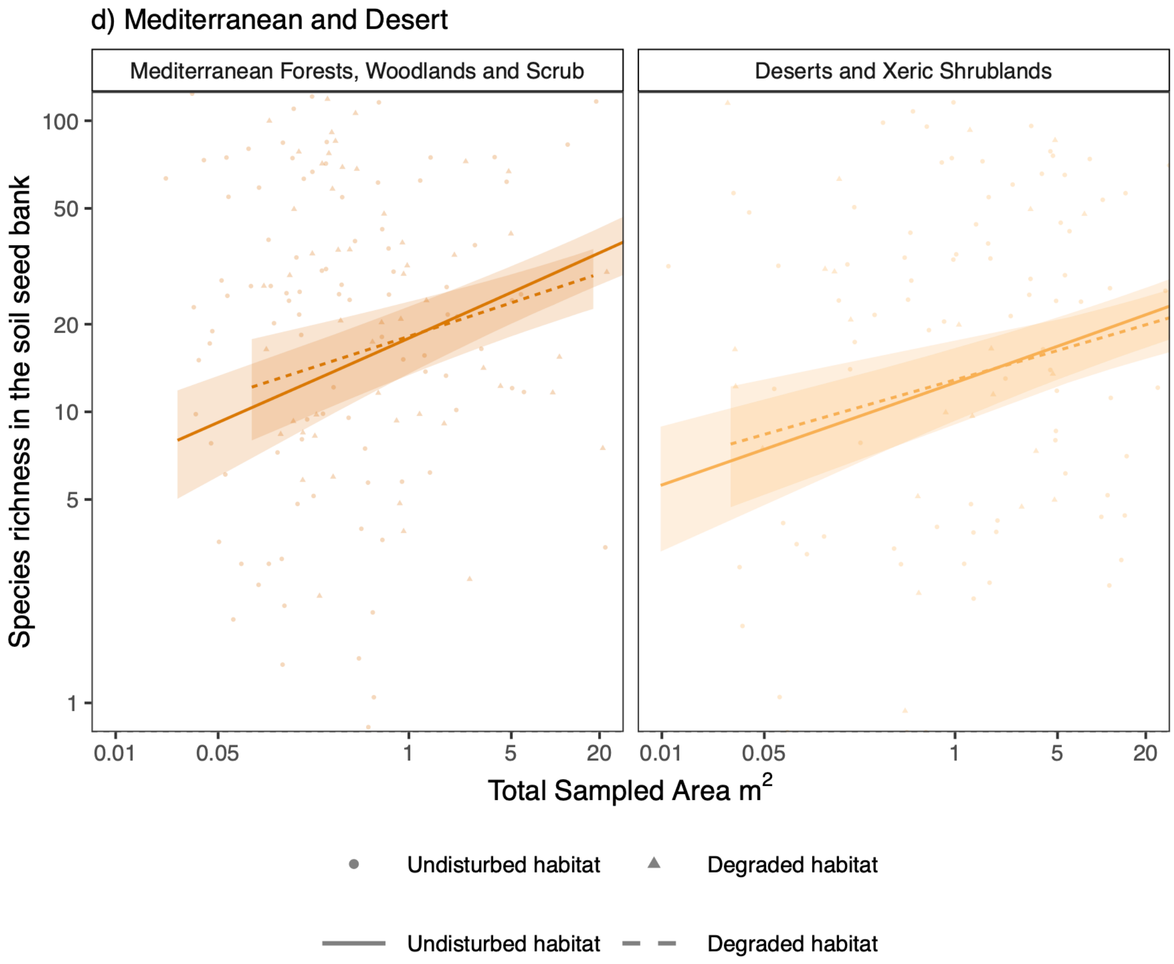


**Figure S2d. Species richness in the soil seed bank as a function of total soil samples ($m^2$).** Each point and line type represents an undisturbed habitat (circles, solid line), and disturbed habitat (triangles, dotted line). Each point is the species richness reported for a given total area of soil sampled in the literature, each line is the overall estimated relationship between the number of species and sampled area for one site, and the shaded area around lines are the 95% credible intervals.

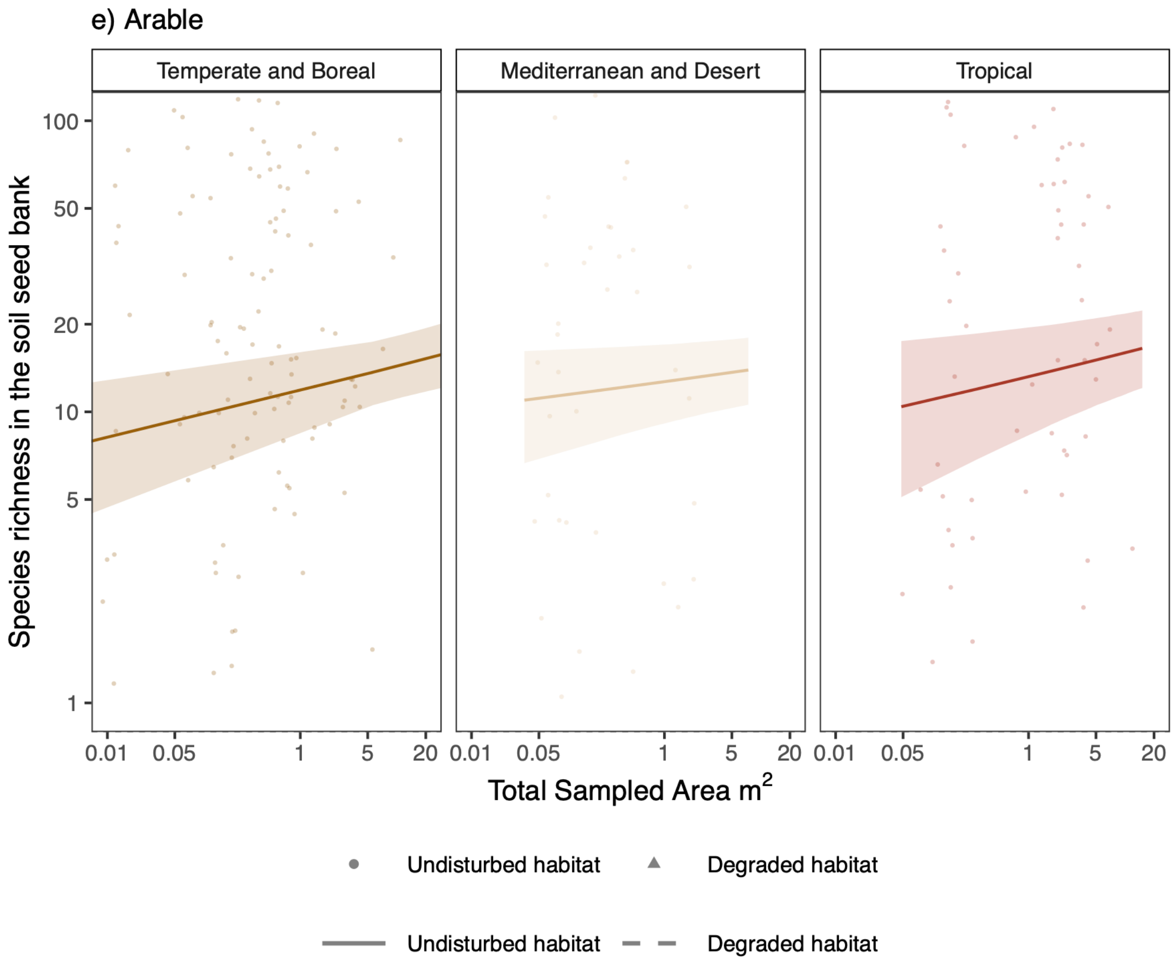


**Figure S2e. Species richness in the soil seed bank as a function of total soil samples ($m^2$).** Each point and line type represents an undisturbed habitat (circles, solid line), and disturbed habitat (triangles, dotted line). Each point is the species richness reported for a given total area of soil sampled in the literature, each line is the overall estimated relationship between the number of species and sampled area for one site, and the shaded area around lines are the 95% credible intervals.

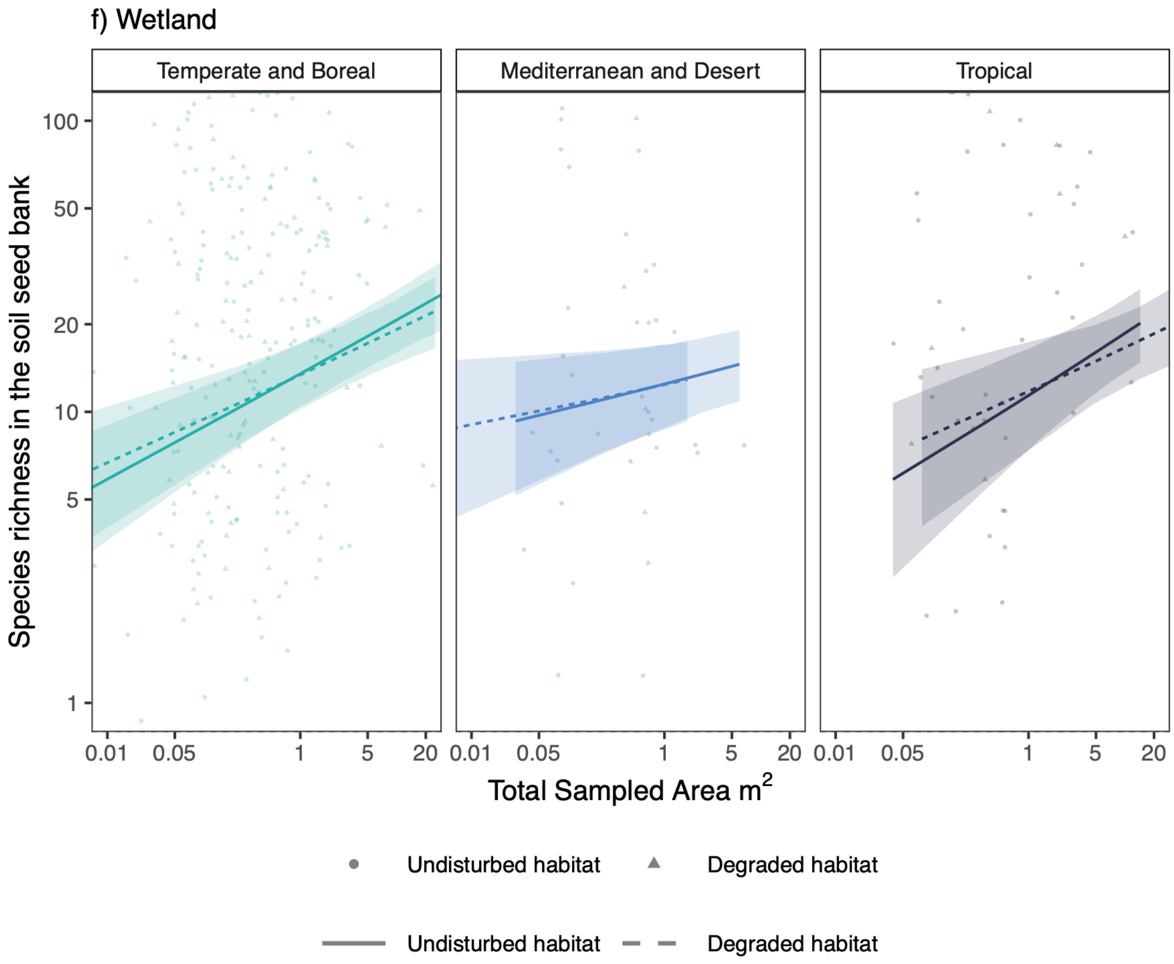


**Figure S2f. Species richness in the soil seed bank as a function of total soil samples ($m^2$).** Each point and line type represents an undisturbed habitat (circles, solid line), and disturbed habitat (triangles, dotted line). Each point is the species richness reported for a given total area of soil sampled in the literature, each line is the overall estimated relationship between the number of species and sampled area for one site, and the shaded area around lines are the 95% credible intervals.

.

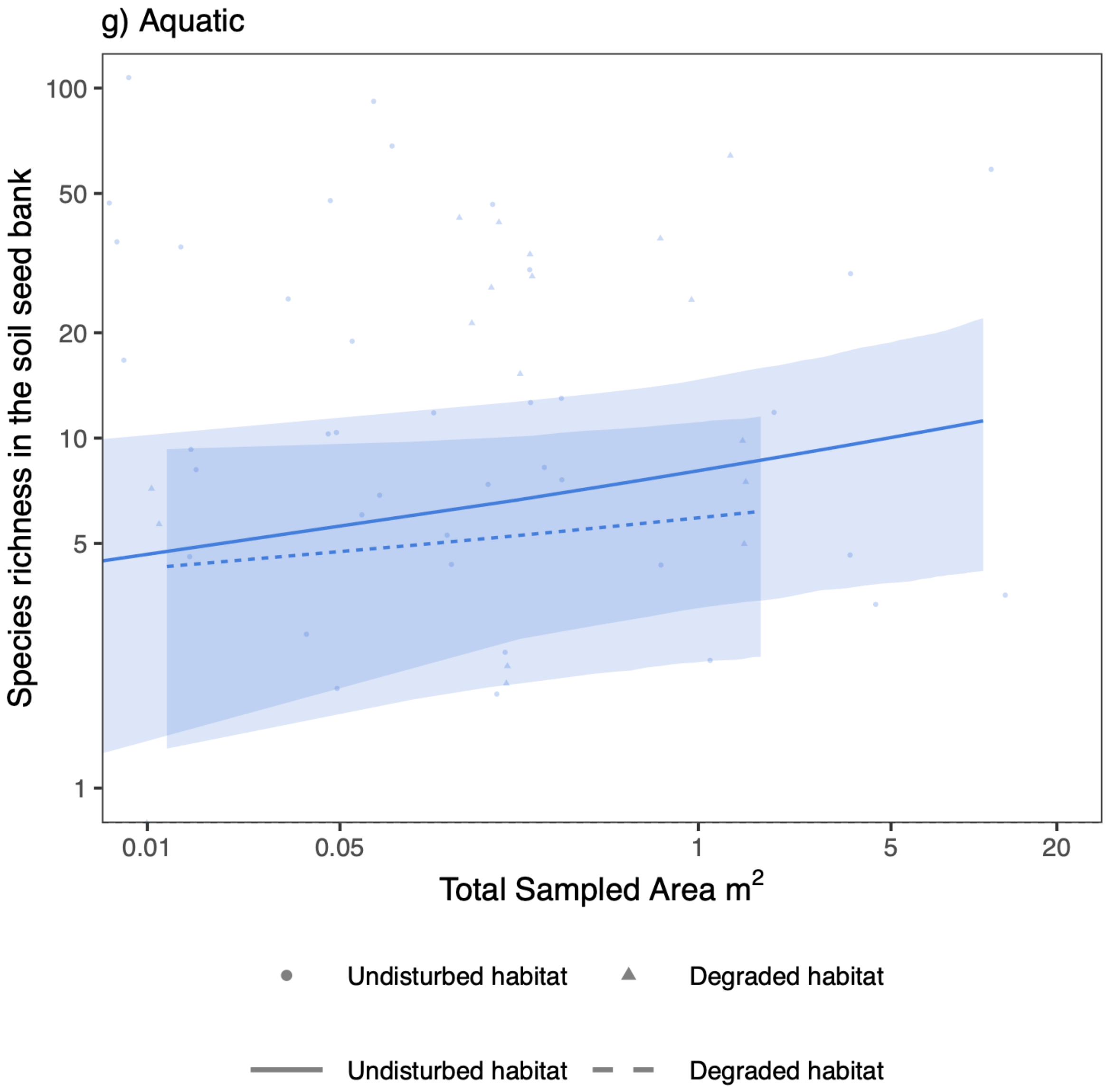


**Figure S2g. Species richness in the soil seed bank as a function of total soil samples ($m^2$).** Each point and line type represents an undisturbed habitat (circles, solid line), and disturbed habitat (triangles, dotted line). Each point is the species richness reported for a given total area of soil sampled in the literature, each line is the overall estimated relationship between the number of species and sampled area for one site, and the shaded area around lines are the 95% credible intervals.
